# A microglia-containing cerebral organoid model to study early life immune challenges

**DOI:** 10.1101/2024.05.24.595814

**Authors:** Alice Buonfiglioli, Raphael Kübler, Roy Missall, Renske De Jong, Stephanie Chan, Verena Haage, Stefan Wendt, Ada J. Lin, Daniele Mattei, Mara Graziani, Brooke Latour, Frederieke Gigase, Haakon B. Nygaard, Philip L. De Jager, Lot D. De Witte

## Abstract

Prenatal infections and activation of the maternal immune system have been proposed to contribute to causing neurodevelopmental disorders (NDDs), chronic conditions often linked to brain abnormalities. Microglia are the resident immune cells of the brain and play a key role in neurodevelopment. Disruption of microglial functions can lead to brain abnormalities and increase the risk of developing NDDs. How the maternal as well as the fetal immune system affect human neurodevelopment and contribute to NDDs remains unclear. An important reason for this knowledge gap is the fact that the impact of exposure to prenatal risk factors has been challenging to study in the human context. Here, we characterized a model of cerebral organoids (CO) with integrated microglia (COiMg). These organoids express typical microglial markers and respond to inflammatory stimuli. The presence of microglia influences cerebral organoid development, including cell density and neural differentiation, and regulates the expression of several ciliated mesenchymal cell markers. Moreover, COiMg and organoids without microglia show similar but also distinct responses to inflammatory stimuli. Additionally, IFN-γ induced significant transcriptional and structural changes in the cerebral organoids, that appear to be regulated by the presence of microglia. Specifically, interferon-gamma (IFN-γ) was found to alter the expression of genes linked to autism. This model provides a valuable tool to study how inflammatory perturbations and microglial presence affect neurodevelopmental processes.

## Introduction

Neurodevelopmental disorders (NDDs), including autism spectrum disorders, intellectual disability and schizophrenia, are chronic, incurable, and unpreventable, imposing a significant burden on affected individuals, their families, and society (Hall et al., 2023; Han et al., 2021). These conditions are caused by aberrant development of the brain, but the underlying molecular and cellular mechanisms are heterogeneous and still hardly understood. This gap in knowledge is hampering the development of novel strategies for prevention and treatment.

Already for a long time, it has been known that pre- and perinatal infections, such as *Toxoplasma Gondii*, rubella virus, or Zikavirus can cause NDDs. Epidemiological studies have also shown a link between NDDs and other infections, such as influenza, although these associations are less clear (Devaraju et al., 2023; Kim et al., 2024; Yates & Mulkey, 2024). Activation of the immune system of the mother during pregnancy, referred to as maternal immune activation (MIA), has been suggested to be a common mechanism explaining the association between prenatal infections and NDDs. A role for MIA in NDDs is further supported by human and animal studies (Brown & Meyer, 2018; Estes & McAllister, 2015; Gumusoglu & Stevens, 2019). Genetic studies on NDDs additionally suggest a connection with immune pathways, as genetic risk factors have been linked to genes that play a role in the immune system (Arenella et al., 2022; Arenella et al., 2023; Iakunchykova et al., 2024). In addition, it is increasingly recognized that microglia, the resident immune cells of the brain (Bilbo & Stevens, 2017), play a crucial role in neurodevelopment and maintaining brain homeostasis (Borst et al., 2021). Microglia have shown to be crucial at shaping the brain at a structural, molecular and functional level, via the regulation of several developmental processes, such as synaptogenesis, synaptic pruning and elimination of neurons (Hammond et al., 2018; Lukens & Eyo, 2022; Wu et al., 2015). Disrupted microglial functions, such as synaptic pruning, can result in structural and functional brain abnormalities, increasing the risk of developing NDDs (Courchesne et al., 2020; Mattei & Notter, 2020; Mondelli et al., 2017). Additionally, animal studies have proposed a role for MIA in disrupting microglial functions in adulthood (Loayza et al., 2023; Traetta & Tremblay, 2022).

How prenatal infections and maternal immune activation affect human neurodevelopment and contribute to NDDs remains unclear. A key reason for this knowledge gap is the difficulty in studying inflammation’s role in a human specific context of developmental diseases, as inflammatory events occur long before birth and diagnosis. Modeling the effects of prenatal risk factors is also challenging and currently relies primarily on exposing rodents to the identified stressors (Meyer, 2023). Given the significant differences between rodent and human neurodevelopment, as well as their response to environmental triggers, we aim to enhance and characterize methods for modeling the impact of prenatal inflammatory stressors on the developing brain using immortalized pluripotent stem cell (iPSC) technology.

Over the past decade, the iPSC field has seen rapid development, and this has resulted in a variety of protocols to explore brain development (Dixon & Muotri, 2023; Lee et al., 2020). Brain organoids are self-organizing, organ-like structures that offer a tool to study the human brain during development (Lancaster et al., 2013) and disease, including NDDs (Mariani et al., 2015; Sarieva et al., 2023). However, they are derived from ectodermal lineage and most models lack other germinal layer-derived cells, such as microglia.

Thus, we aim to further develop and characterize brain organoids to study immune processes and model maternal immune activation by exposing them to inflammatory insults. We used an adapted model following published protocols (Fagerlund et al., 2021; Park et al., 2023), to generate cerebral organoids with microglia (COiMg). We first characterized the effect of the presence of microglia on the cerebral organoids, and subsequently analyzed how these organoids respond to a panel of inflammatory stimuli, including Lipopolysaccharide (LPS), Interferon (IFN)-α, IFN-γ and interleukin (IL)-6. Finally, we focused on how the cytokine IFN-γ, influences cerebral organoid development at both transcriptional and protein levels, and the role microglia play in this process.

The present study provides a valuable platform for modeling the developing brain and for investigating the role of microglia when exposed to immune challenges.

## Results

### Characterization of a microglia-containing cerebral organoid model

To study the impact of inflammatory insults on the developing human brain, it is important to use a human model that reflects developing human brain tissue and includes microglia. Microglia are derived from myeloid progenitor cells that develop in the yolk sac, and that enter the developing brain in very early stages. In an attempt to mimic this developmental trajectory, we used iPSC-derived hematopoietic stem cells (HPCs) and added these cells to cerebral organoids that had been in culture for 25-28 days, as illustrated in the schematic protocol in Figure 1A. We used HPCs that constitutively express GFP to track these cells in the organoids. After 7 days of co-culture, we found GFP-expressing cells throughout the organoids while the overall structural composition of the organoids was not affected (Fig.1B). GFP-positive cells exhibited an amoeboid morphology with an initial onset of ramification, consistent with previous findings in organoids (Fagerlund et al., 2021; Park et al., 2023), and these cells expressed the myeloid/microglia markers IBA1 and CD45 (Fig.1C,D). By using flow cytometry (gating strategy showed in Fig.S1A), we found that between 0.5% and 4% of the cells in the organoids expressed GFP (Fig.1E,F). GFP^+^ cells express high levels of CD45, and CD11b, P2RY12, and TREM2 (Fig.1G), while expression of CD14 and CD206 was low or absent (Fig.S1B). Exposure of COiMg, but not CO, to LPS, resulted in the release of TNF-α and IL-8 (Fig.1G,I).

**Figure 1.**
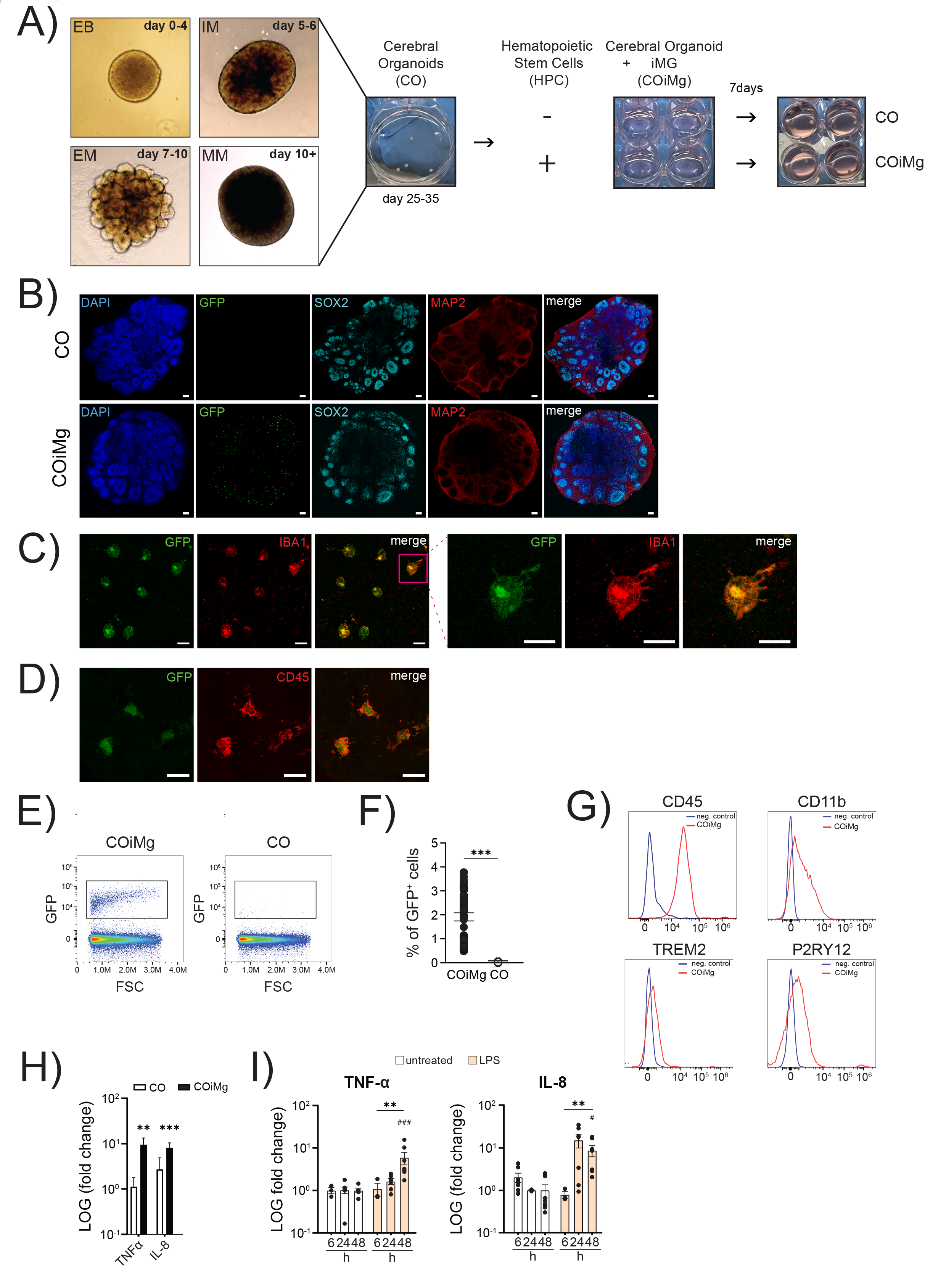
Generation and characterization of an immune-competent cerebral organoid model that responds to inflammatory triggers. (A) Schematic illustration of the generation of COiMg, including CO differentiation and Cerebral Organoid differentiation and the co-culturing protocol. (B) Representative images of CO and COiMg stained with the neural markers SOX2 and MAP2. Microglia are labelled with GFP. HOECHST was used for nuclei staining. 10X Magnification. Scale bar, 100µm. (C,D) Representative images of COiMg stained with the microglial markers IBA1 (C) and CD45 (D). Microglia are labelled with GFP. 20X and 63X Magnification. Scale bar, 20 and 10 µm, respectively. (E) CO (in white) and COiMg (in black) were dissociated and FACS was performed to analyze the presence of microglia, expressed as GFP^+^ cells. Density plot of COiMg and CO gated for GFP^+^ cells. Cells were pre-gated for singlets and viability. (F) Percentages of GFP^+^ cells were calculated out of single viable cells in both CO and COiMg. Data are represented as points and error bars as mean ± SEM. N= 3 biological replicates (batches). Welch’s t-test was performed. *** P ≤ 0.001. (G) Histogram plots depicting the microglial marker CD45, CD11b, TREM2 and P2RY12 expression in COiMg (in red). Unstained sample was used as negative control. (H) CO (in white) and COiMg (in black), derived from MSN38 iPSC donor line, were stimulated with LPS (100 ng/mL) for 48 h. TNF-α and IL-8 release was determined via ELISA and expressed as fold change release normalized to untreated samples. Mann Whitney t-test was performed. ** P ≤ 0.01, *** P ≤ 0.001. Data are represented as bars and error bars as mean ± SEM. N= 3 or more biological replicates (batches). (I) COiMg were stimulated with LPS (100 ng/mL) for 6, 24 and 48 h. TNF-α and IL-8 release was determined via ELISA and expressed as fold change release normalized to untreated samples. Kruskal-Wallis test, followed by Dunn’s post-hoc test, was performed. # P ≤ 0.01, ** P ≤ 0.01, ### P ≤ 0.001. Data are represented as mean ± SEM. N= 3 or more biological replicates (batches).

### COiMg display microglial gene signatures

To further characterize the COiMg model, we performed bulk RNA sequencing on three different donor lines. Principal Component Analysis (PCA) revealed a clear separation between CO and COiMg (Fig.2A). Differential Expressed Gene (DEG) analysis resulted in 283 significantly upregulated and 209 significantly downregulated genes (Fig.2B). Clustering based on a panel of well-characterized microglia genes showed a clear separation between CO and COiMg (Fig.2C). Among the upregulated genes the top 20 DEGs included markers related to microglial ontogeny such as *SSP1*, *CX3CR1*, *IRF8*, *CSF1R*, pattern recognition receptor genes including *TLR2*, *4* and *6*, as well as genes related to synaptic remodeling and phagocytosis such as *C3*, *C1Q*, *CD68*, *TREM2*, *TYROBP* and *SYK* (Fig.2D,E,F). To further confirm the expression of microglia markers in COiMg, we compared the expression of *AIF1, CX3CR1, TREM2, P2RY12* and *TMEM119* between CO and COiMg, as well as organoid-conditioned microglia (ocMG), primary microglia isolated from post-mortem human brain tissue (pMG) and iPSC-derived microglia (iMG) (Fig.2SA,B). *AIF1* showed significant upregulation in COiMg compared to CO. All microglial markers were expressed in COiMg. As microglia are only a minority of the cells in the organoids, the expression was lower in COiMg compared to the monocultures. Gene Ontology (GO) analysis (Fig.S2C) confirmed that differentially expressed genes were significantly enriched in immune response pathways, including phagocytosis, immune receptor activity, neuroinflammation, MHC I and microglial identity. Conversely, downregulated genes were enriched in pathways related to blood circulation, cilia movement and extracellular matrix (Fig.S2D). We performed single-cell RNA sequencing (scRNAseq) on CO and COiMg from two different donor lines. We analyzed 107,420 cells in total (Fig.2G) and identified 8 distinct clusters, which mapped to four major cell-types. We found one cluster with a microglial signature including *AIF1, C1QA, C1QB, CSF1R, CX3CR1, GPR34, ITGAM, P2RY12, TREM2, TMEM119* in the COiMg specifically.

**Figure 2.**
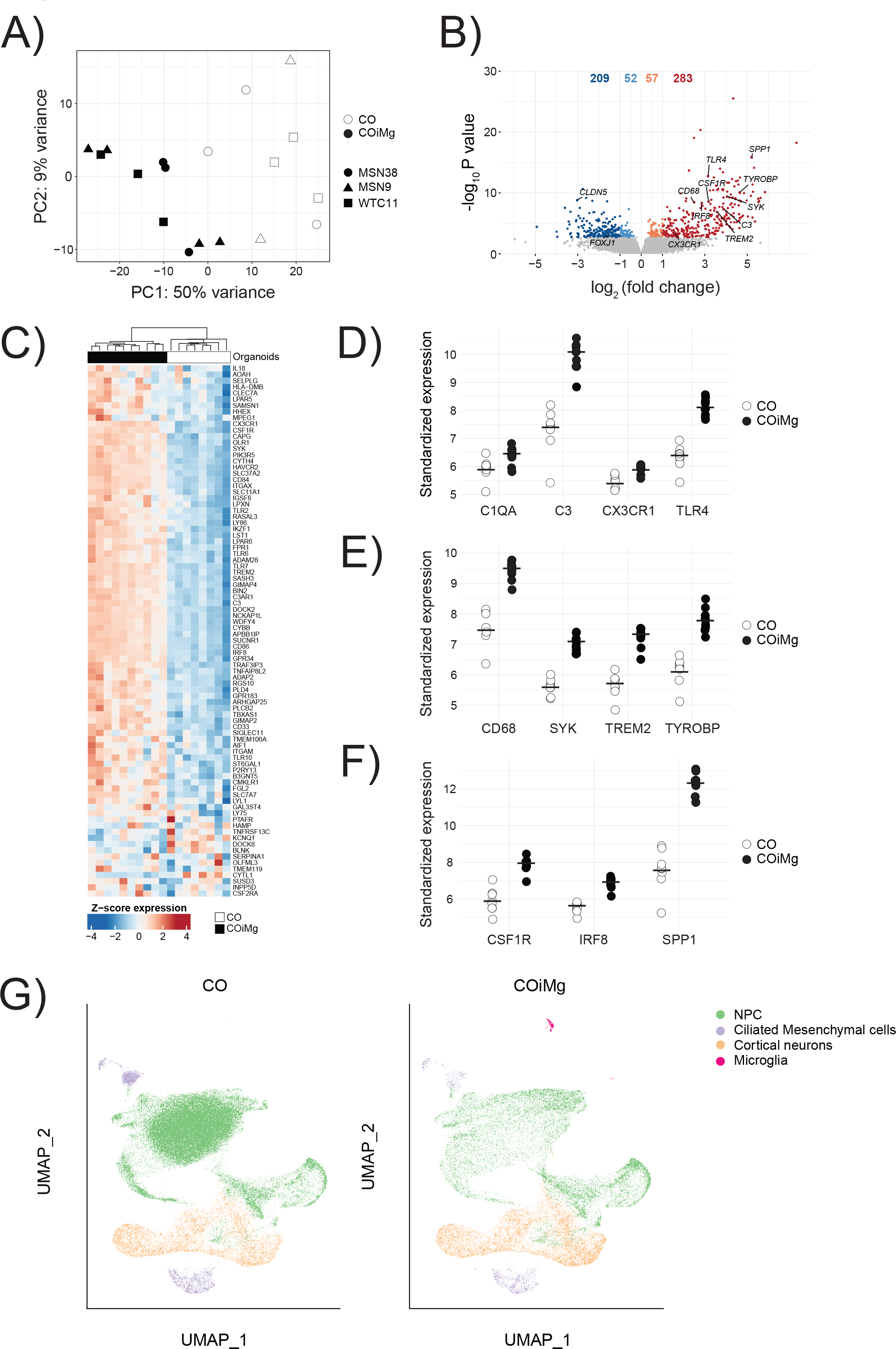
Microglia-containing cerebral organoids express microglial genes. (A) Bulk RNAseq PCA Plot of the first two PCs displaying CO and COiMg sample distance in all three donor iPSC lines. N=8-9 per condition, and N=3 per line (white = CO, black = COiMg, circle = MSN38, triangle=MSN9, square = WTC11). PCs were calculated based on the top 500 variable genes. (B) Volcano plot showing differentially expressed genes. Microglial genes were annotated. Each dot represents a gene. Colorization shows genes differentially expressed at fold change < 0.1 (lightblue = downregulated at log2FC < 0, orange = upregulated at log2FC > 0, darkred = upregulated at log2FC > 1, darkblue = downregulated at log2FC < -1). (C) Heatmap of row-scaled vst-standardized and corrected gene expression of microglia genes in CO and COiMg. (D,E,F) Dotplots of vst-standardized and corrected expression of specific microglial markers: *C1QA, C3, CX3CR1, TLR4, CD68, SYK, TREM2, TYROBP, CSF1R, IRF8 and SPP1*. Each dot represents one organoid. (G) UMAP of CO and COiMg showing cluster identity for NPC, Ciliated Mesenchymal cells, Cortical Neurons and Microglia. Each dot is a single cell.

The impact of the microglial co-culture on organoid structure and developmental trajectories of neurons.

As microglia have been shown to be involved in neurogenesis and phagocytosis of synapses (Cunningham et al., 2013), we investigated the microglial contribution to the cerebral organoid structure by using immunofluorescence staining (Fig.3A). We first analyzed the effect of microglia on cell density. HOECHST staining revealed a similar trend for the three different lines (Fig.S3A), with a significantly decreased cell density in COiMg when analyzed all lines together (Fig.3B). Next, we analyzed the impact of microglia on the number of rosettes (Fig.3C and Fig.S3B,C), typical structures which are thought to resemble processes related to neural tube development and foci of NPC development (Hong et al., 2023). We found a significantly decreased number of rosettes in COiMg compared to CO when all lines were analyzed together (Fig.3C), but the results were variable between the lines (Fig.S3C). To further evaluate the impact of microglia on neuronal composition, we performed immunofluorescence staining for the neuro- progenitor (SOX2) and mature neuron (NEUN) markers. We observed no significant changes in the percentage of SOX2^+^ cells (Fig.3D), but there was a significant decrease in the proportion of NEUN^+^ cells in COiMg compared to CO (Fig.3E), despite variability across organoids and cell lines (Fig.S3D,E).

**Figure 3.**
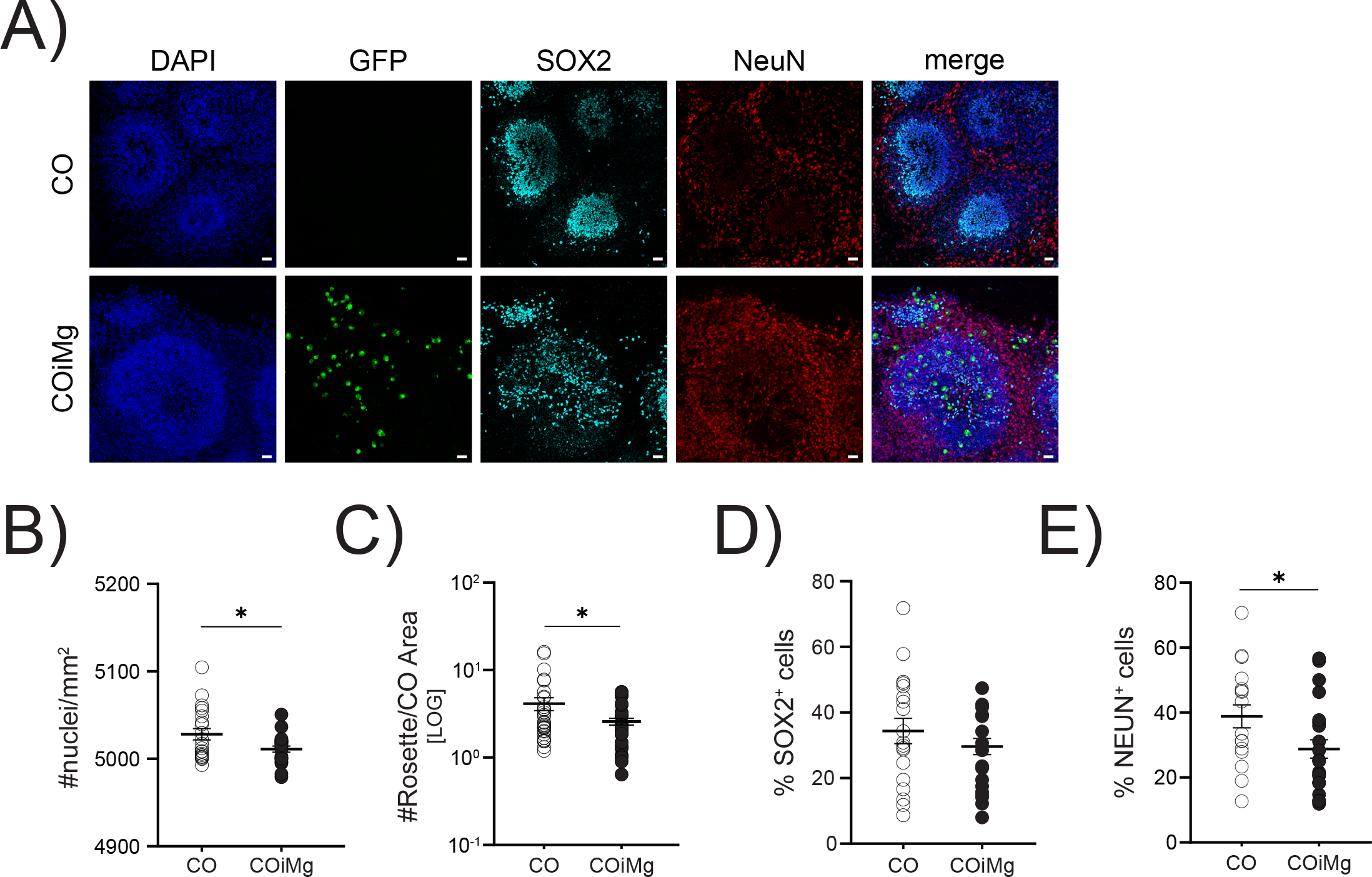
Microglia presence decreased cell density and the percentage of NEUN^+^ population in cerebral organoids. (A) Representative images of CO and COiMg stained with the neural stem cell marker SOX2 and the neuronal marker NEUN. Microglia are labelled with GFP. HOECHST was used for nuclei staining. 20X Magnification. Scale bar, 10µm. (B) Quantification of number of nuclei per mm^2^ was performed in CO and COiMg. Data are represented as points and error bars as mean ± SEM. N= 3 biological replicates (batches) per line. Welch’s t-test was performed. * P ≤ 0.05. (C) Quantification of number of rosettes per organoid area was performed in CO and COiMg. Data are represented as points and error bars as mean ± SEM. N= 3 biological replicates (batches) per line. Welch’s t-test was performed. * P ≤ 0.05. (D,E) Percentage of SOX2^+^ (D) and NeuN^+^ (E) cells was calculated in CO and COiMg. Data are represented as points and error bars as mean ± SEM. N= 3 biological replicates (batches) per line; all three lines were pulled. Unpaired t-test was performed. * P ≤ 0.05.

Microglia are an important source of TNF-α, and, given TNF-α’s known role in neurogenesis (Iosif et al., 2006; Mattei et al., 2014) through its interaction with both the pro-apoptotic TNFR1 and the proliferative TNFR2 (Keohane et al., 2010), we sought to investigate whether mature neurons favor one receptor over the other, potentially elucidating the reduced NEUN^+^ cell count in COiMg. Our scRNAseq revealed a notable upregulation of the pro-apoptotic receptor TNFR1 in cortical neurons of COiMg compared to CO (Fig.S3F). This observation could imply the involvement of an apoptotic pathway as a potential mechanism for neuronal reduction. Subsequently, both CO and COiMg were functionally tested in terms of neuronal firing. Cerebral organoids were plated in 48- well multi electrode array (MEA) plates for the monitoring of electrical activity with or without microglia. Both CO and COiMg displayed single spike firing and burst-like activity (Fig.S3G). Although COiMg tended to attach better to the surface of MEA plates, resulting in overall higher activities, the overall firing rate did not change in the presence of microglia (Fig.S3H). Application of 1µM Tetrodotoxin (TTX) to block any action potentials from firing, effectively blocked all electrical activity from all organoids, resulting in a significant drop in burst frequencies (Fig.S3I). As a result, we conclude that neurons in cerebral organoids, with or without microglia, display functional activity, including single cell firing as well as coordinated burst firing, with no significant difference between the two conditions.

### Microglia influence the expression of cilia-related genes

As previously mentioned, the presence of microglia appears to downregulate pathways involved in ciliated mesenchymal cell-related genes (Supplementary Table 5). Ciliated mesenchymal cells have been reported to regulate the formation of the choroid plexus (ChP), a brain structure responsible for producing the cerebrospinal fluid (CSF), which is important for the development and maintenance of the brain (Lehtinen et al., 2013; Lun et al., 2015; Redzic et al., 2005) and vascularization (Pombero et al., 2016). In both sc- and bulk RNA sequencing analyses, we observed downregulation of transcripts, including Forkhead Box J1 (*FOXJ1*), Claudin 5 (*CLDN5*) and Transthyretin (*TTR*) (Fig.4A,B and Fig.S4A). FOXJ1 is a transcription factor that regulates the development and maturation of ciliated cells, such as radial glia differentiating into ependymal cells (Jacquet et al., 2009; Yu et al., 2008), which are necessary for the formation of ventricular structures such as the choroid plexus. ScRNAseq confirmed that FOXJ1 is predominantly expressed by mesenchymal cells and to a lesser extent by NPCs (Fig.S4B). Both sc- and bulk RNA sequencing revealed downregulation of FOXJ1 in COiMg (Fig.4A,B and Fig.S4A). Additionally, we validated FOXJ1 protein expression using immunofluorescence, which showed a decreased percentage of FOXJ1^+^ cells in COiMg compared to CO (Fig.4C,D and Fig.S4C). CLDN5 belongs to the family of claudins, proteins important for the formation of tight junctions essential for cell-cell communication and brain vasculature, including the brain blood barrier (Günzel & Yu, 2013). CLDN5 is primarily expressed by endothelial cells, and it is the most abundant claudin in the brain (Garcia et al., 2022; Vanlandewijck et al., 2018). Our scRNAseq indicated that mesenchymal cells are the primary source of CLDN5, with contributions from NPCs and cortical neurons to a lesser extent, confirming its importance as the most expressed claudin in the brain (Fig.S4B). Both sc- and bulk RNAseq analysis revealed downregulation of CLDN5 in COiMg compared to CO, with the transcript downregulation primarily originating from mesenchymal and NPCs (Fig.4A,B and Fig.S4A). Immunofluorescence confirmed lower CLDN5 expression in COiMg compared to CO (Fig.4E,F and Fig.S4D). Finally, TTR is a protein expressed by endothelial cells of the choroid plexus and is involved in the transport of thyroxine and retinol into the CSF (Herbert et al., 1986). Sc-RNAseq revealed that TTR is abundantly expressed by mesenchymal cells (Fig.S4B), but the transcriptomic downregulation is exclusively observed in NPCs and neuronal cells (Fig.4B and Fig.S4A). Further validation of TTR protein expression in the organoid tissue showed a decreased signal in COiMg compared to CO (Fig.4G,H and Fig.S4E). Additionally, CLDN5 and TTR were found to colocalize, as shown in Figure S4F. In conclusion, COiMg display reduced RNA and protein expression of specific markers enriched in ciliated mesenchymal cells.

**Figure 4.**
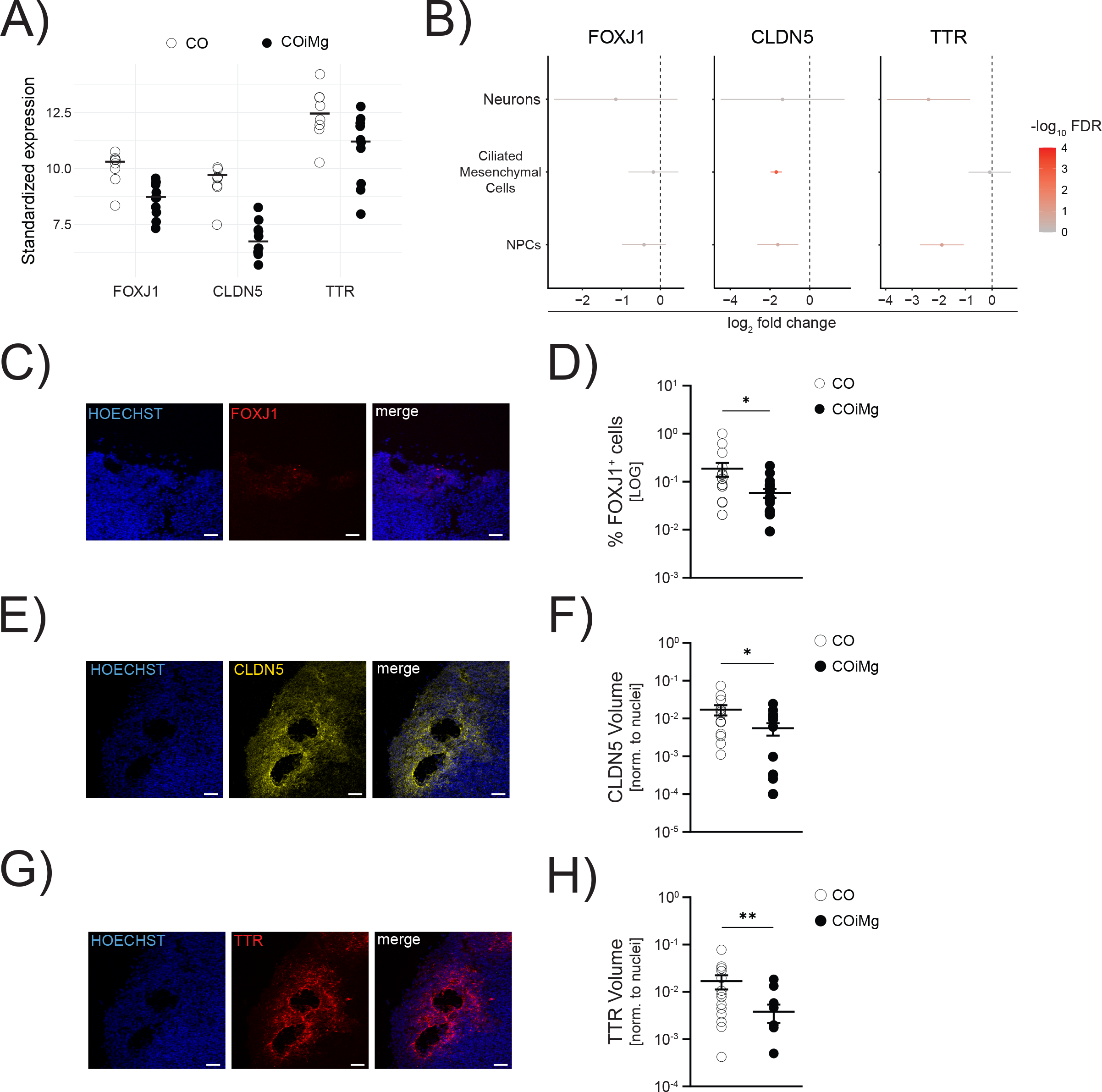
Microglia presence decreases the expression of cilia-associated markers on the transcriptional and protein level. (A) Dotplots of vst-standardized and corrected expression of *CLDN5*, *FOXJ1* and *TTR* genes derived from bulk RNAseq. Each dot represents one organoid. (B) Downregulated expression of *CLDN5*, *FOXJ1* and *TTR* in Neurons, Ciliated Mesenchymal Cells and Radial Glia, extracted from the scRNAseq. (C) Representative image of CO stained with FOXJ1. HOECHST was used for nuclei staining. 20X Magnification. Scale bar, 50µm. (D) Percentage of FOXJ1^+^ cells quantified in CO and COiMg. Data are represented as points and error bars as mean ± SEM. N= 2 biological replicates (batches) per line. Mann Whitney t-test was performed. * P ≤ 0.05. (E) Representative image of CO stained with CLDN5. HOECHST was used for nuclei staining. 20X Magnification. Scale bar, 50µm. (F) Quantification of CLDN5 surface volume normalized to HOECHST volume was performed in CO and COiMg. Data are represented as points and error bars as mean ± SEM. N= 2 biological replicates (batches) per line. Unpaired t-test was performed. * P ≤ 0.05. (G) Representative image of CO stained with TTR. HOECHST was used for nuclei staining. 20X Magnification. Scale bar, 50µm. (H) Quantification of TTR surface volume normalized to HOECHST volume was performed in CO and COiMg. Data are represented as points and error bars as mean ± SEM. N= 2 biological replicates (batches) per line. Mann Whitney t-test was performed. ** P ≤ 0.01.

### COiMg respond to inflammatory stimuli

Using CO and COiMg derived from three different control iPSC donor lines (Fig.5A), we sought to determine the capacity of organoids to respond to inflammatory stimuli. CO and COiMg were challenged with LPS, IFN-γ, IFN-α and IL-6 for 24 and 48 hours and, subsequently, supernatants were collected and analyzed for cytokine levels using multiplex ELISA. We observed different behaviors based on the stimulant but also on the inflammatory molecule released. We focused on LPS as well as the type I and II IFNs due to their strong effect. Overall, we observed that the response to LPS intensified over time, while the response to IFN-γ and IFN-α remained consistent, suggesting a potential saturation of the signal (Fig.S5A). When exposed to LPS, COiMg, but not CO, released TNF-α, IL-8, IL-10, CXCL9, IL-6, MCP-1, and MCP-3, suggesting a microglia-dependent response (Fig.5B and Fig.S5B). IFN-γ response appeared to be independent on the microglia presence as COiMg and CO did not differ in terms of response to the selected inflammatory molecules analyzed. However, CO exhibited higher CXCL10 production compared to COiMg (Fig.5C and Fig.S5C). The response to IFN-α varied; while some cytokines showed a microglia-dependent response similar to LPS, others mirrored IFN-γ responses. Specifically, TNF-α, IL-6 and IL-10 release was depending on microglia presence, while CXCL10 release did not differ between CO and COiMg (Fig.5D and Fig.S5D). We performed the same ELISA immunoassay with supernatants from stimulated ocMG and iMG and found similarities between the responses of COiMg and these microglial monocultures (Fig.5E). We expanded our analysis by conducting bulk RNAseq on untreated and 48-hour LPS- and IFN-γ-treated CO and COiMg. Although significant changes were observed with IFN-γ, we found no significant transcriptional differences when organoids were stimulated with LPS compared to untreated samples (Fig.5F), suggesting that LPS may require a chronic exposure for changes or different time windows for analysis. Moreover, a clear separation between IFN-y stimulated CO and COiMg transcriptomic profiles was visible.

**Figure 5.**
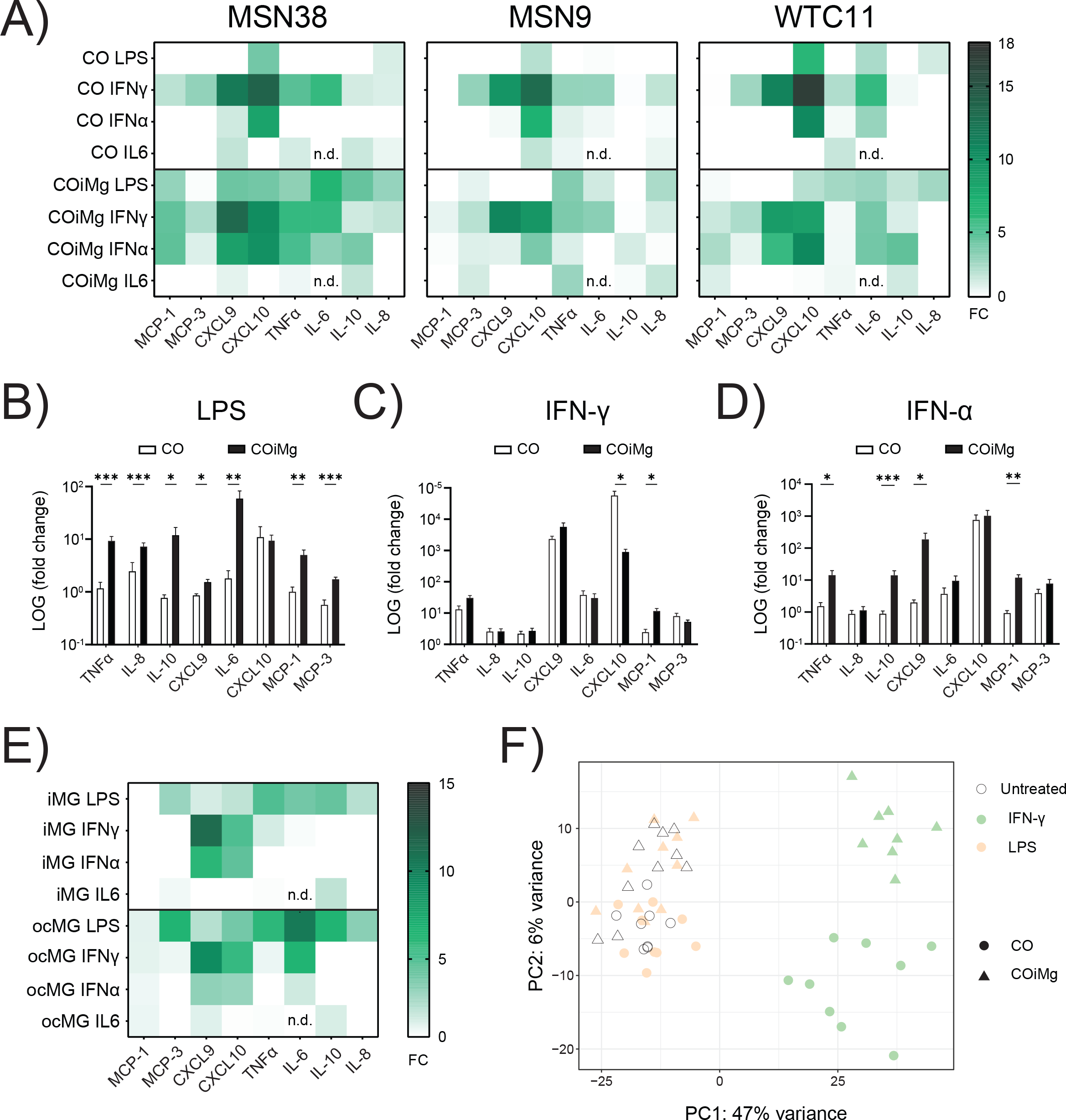
Effect of different inflammatory triggers on cerebral organoids with or without microglia. (A) CO and COiMg, derived from all three donor iPSC lines, were stimulated for 48 h with the inflammatory ligands LPS, IFN-γ, IFN-α and IL6 (all at a concentration of 100 ng/mL), supernatants were collected, and cytokine and chemokine release was analyzed via Multiplex ELISA. Data are shown in a heatmap representing cytokine release normalized to untreated conditions. N= 3 biological replicates (batches) per line. (B,C,D) CO and COiMg were stimulated for 48 h with the inflammatory ligands LPS (B), IFN-γ (C) and IFN-α (D), supernatants were collected and cytokine and chemokine release was analyzed via Multiplex ELISA. Data, expressed as fold change release compared to untreated conditions, are shown as bars and error bars as mean ± SEM. N= 3 biological replicates (batches) per line; all three lines were pulled. (E) iMG and ocMG were stimulated for 48 h with the inflammatory ligands LPS, IFN-γ, IFN-α and IL6, supernatants were collected and cytokine and chemokine release was analyzed via Multiplex ELISA. Data are shown in a heatmap representing cytokine release normalized to untreated conditions. N= 3 per microglial model. (F) PCA plot of the first two principal components displaying CO and COiMg usample distances in untreated and stimulated conditions (white = untreated, orange = LPS and lightgreen = IFN-γ) from bulk RNAseq. N= 3 per stimulants per iPSC line. PCs were calculated based on the top 500 variable genes.

### Exposure to IFN-γ induces significant transcriptional changes in cerebral organoids

To study the effect of inflammatory insults on the organoid transcriptome, CO and COiMg, derived from three donor lines, were stimulated with IFN-γ for 48 hours, and the tissue was dissociated and processed for bulk RNAseq. PCA analysis of gene expression showed a clear separation between untreated and stimulated in both CO and COiMg samples (Fig.6A and Fig.S6A), with no transcriptional differences observed between the three iPSC donor lines.Among the top upregulated genes in both IFN-γ-treated CO and COiMg, we identified Chemokine Ligand 9 and 10 (*CXCL9* and *CXCL10*), as well as Indoleamine 2,3 dioxygenase 1 (*IDO1*) and Guanylate binding protein (*GBP5*) (Fig.6B and Fig.S6B,C,D). CXCL9 and CXCL10 are chemokines involved in the innate and adaptive response to IFN-γ by attracting and recruiting NK and T cells (Ellis et al., 2010). ELISA analysis confirmed the release of CXCL9 and CXCL10 at the protein level when the organoids were exposed to IFN-γ (Fig.5A,C and Fig.S5A,C), consistent with previous reports (Rock et al., 2005). IDO1 is an enzyme responsible for the degradation of the essential amino acid L-Tryptophan and is involved in several biological processes, including immune homeostasis (Pallotta et al., 2022). GBP5 belongs to the GTPase subfamily and is involved in inflammasome activation and innate immune response against pathogens (Shenoy et al., 2012). Both IDO1 and GBP5 expression and activity have been shown to be induced by IFN-γ. Immunofluorescence microscopy confirmed the increased expression of IDO1 when CO and COiMg were treated with IFN-γ (Fig.6C,D and Fig.S7A,B). IDO1 is also expressed by innate immune cells in the brain (Kwidzinski & Bechmann, 2007). We found that IFN-γ-treated CO expressed high levels of IDO1 and that approximately 50% of the microglia expressed IDO1 (Fig.6E and Fig.S7C) in COiMg suggesting that other cells types aside microglia are responsible for its expression. Increased GBP5 protein expression was confirmed by immunofluorescence after IFN-γ exposure (Fig.S7D). Given the fact that both CO and COiMg respond to IFN-γ, we used our scRNAseq analysis to determine which cells express the IFN-γ receptor, which is an heterodimer composed of IFNGR1 and IFNGR2 (Bach et al., 1997). By using our scRNAseq analysis we found that while in our COiMg model microglia are the main source of both receptors, mesenchymal and neuronal cells express both receptors in lower amount in both CO and COiMg (Fig.S7E). We next investigated the effect of IFN-γ on the organoid structure and neuronal population. We analyzed nuclei and rosette numbers after IFN-γ stimulation and observed a significant increase in nuclei in COiMg when treated with IFN-γ across all the three iPSC donor lines (Fig.6F and Fig.S8A). However, no changes in the number of rosettes were observed between iPSC donor lines (Fig.S8B,C). Interestingly, treatment with IFN-γ resulted in a significant increase in the overall percentage of microglia (Fig.6G), although we observed variability across lines (Fig.S8D). When we looked at the neuro-progenitor percentage, we observed a significant decrease of SOX2^+^ cells in CO when treated with IFN-γ, while in COiMg no significant changes were detected (Fig.6H), however, we observed line-to-line variability (Fig.S8E). Furthermore, while adding microglia decreased the number of mature neurons, IFN-γ further decreased NEUN^+^ cells in both CO and COiMg (Fig.6I and Fig.S8F). Of note, as shown before for the CO *versus* COiMg comparison, we detected high organoid-organoid and line-line variability for the neuronal analyses. We observed 847 upregulated and 74 downregulated genes when COiMg were exposed to IFN-γ, while we detected 452 upregulated and 189 downregulated genes in CO (Fig.6J,K). Upregulated genes were enriched for microglia, phagocytic, MHC Class II, and neuroinflammatory pathways (Fig.S9A,B). While the upregulated genes related to biological process appear similar in both CO and COiMg, the intensity of expression, indicated in -log(q values), is different (Fig.S10A,B,C,D,E,F).

**Figure 6.**
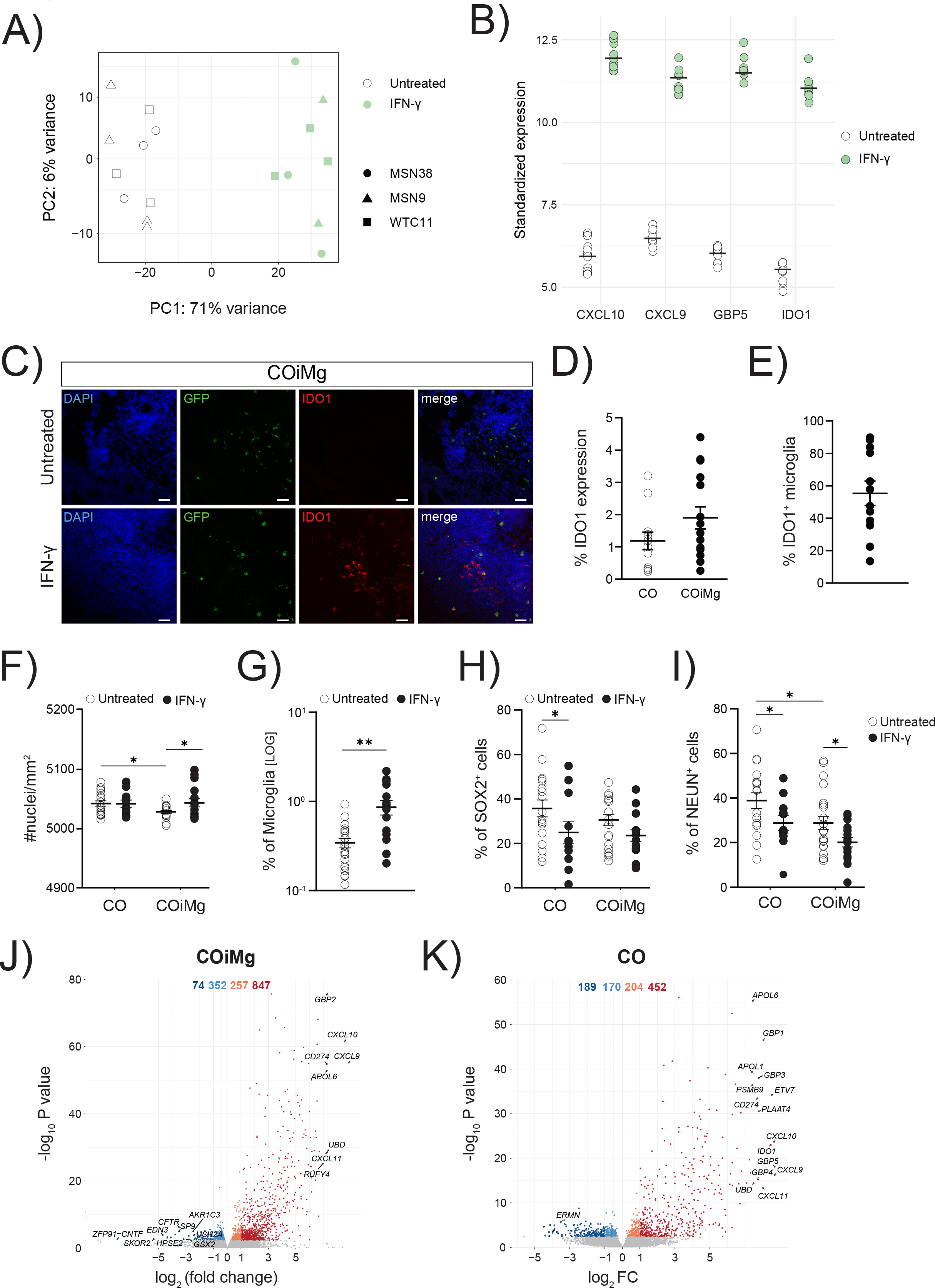
IFN-γ-mediated signaling pathways are upregulated in both CO and COiMg. (A) PCA plot of the first two PCs displaying COiMg sample distances in untreated and stimulated conditions (white = untreated, green = IFN-γ) from bulk RNAseq. N= 3 per stimulants per iPSC line. PCs were calculated based on the top 500 variable genes. (B) Dotplots of vst-standardized and corrected bulk RNAseq expression of the differentially upregulated genes of CXCL10, CXCL9, GBP5 and IDO1 in IFN-γ-treated COiMg. (C) Representative image of COiMg untreated (upper) or treated with IFN-γ (lower) stained with IDO1. Microglia are labelled with GFP. HOECHST was used for nuclei staining. 20X Magnification. Scale bar, 50 µm. (D) Percentage of IDO1 expression in CO and COiMg. Data are represented as points and error bars as mean ± SEM. N= 2 biological replicates (batches) per line. All three lines pulled. (E) Percentage of IDO1^+^ microglia was calculated in COiMg. Data are represented as points and error bars as mean ± SEM. N= 2 biological replicates (batches) per line. All three lines pulled. (F) Quantification of number of nuclei per mm^2^ was performed in untreated and IFN-γ- treated CO and COiMg. Data are represented as points and error bars as mean ± SEM. N= 3 biological replicates (batches) per line. Two-way ANOVA test was performed. * P ≤ 0.05. (G) Percentage of microglia, expressed on a logarithmic scale, was calculated in untreated and IFN-γ-treated COiMg. Data are shown as points with mean ± SEM. N= 2 biological replicates (batches) per line. All three lines pulled. Mann Whitney t-test was performed. ** P ≤ 0.01. (H,I) Percentage of SOX2^+^ (H) and NeuN^+^ (I) cells was calculated in untreated and IFN-γ-treated CO and COiMg. Data are represented as points and error bars as mean ± SEM. N= 3 biological replicates (batches) per line; all donor lines are pulled. Two-way ANOVA test, followed by Fisher’s LSD post-hoc test, was performed. * P ≤ 0.05. (J,K) Volcano plots showing differentially expressed genes in IFN-γ-treated COiMg (J) and CO (K). Each dot represents a gene. Colorization shows genes differentially expressed at fold change < 0.1 (lightblue = downregulated at log2FC < 0, orange = upregulated at log2FC > 0, darkred = upregulated at log2FC > 1, darkblue = downregulated at log2FC < -1).

### Dissecting transcriptional changes of CO and COiMg in response to IFN-γ

Increased levels of IFN-γ during pregnancy have been associated with the risk for autism and schizophrenia (Majerczyk et al., 2022). For this reason, we compared our gene signature with genes that have been associated with the risk of developing different NDDs. We found that the downregulated genes following exposure of cerebral organoids to IFN-γ are significantly linked to rare variants of autism spectrum disorder (ASD) (Fig.7A). The presence of microglia appears to modulate the decreased expression of certain genes by reverting their phenotype or behavior. For instance, Insulin-like growth factor-1 (IGF-1) is a hormone that plays crucial role in pre- and post-natal brain development, myelination, synapse formation, neurotransmitter production, cognition and adult neurogenesis (Nieto-Estévez et al., 2016; Wrigley et al., 2017). IGF-1 has been implicated in and found to be deregulated in ASD (Chen et al., 2014) and is currently being investigated as a potential treatment for syndromic forms of ASD (Costales & Kolevzon, 2016; Vahdatpour et al., 2016). IGF-1 is downregulated in IFN-γ-treated CO compared to untreated ones, while in COiMg no difference is observed (Fig.7B), suggesting a potential protective role of microglia. Two other genes, KCNH5 and UNC5D, which are associated with synaptic functions, exhibited similar behavior as IGF-1, with a downregulation after IFN-y exposure of CO, but not COiMg (Fig.7B). In addition, we observed that the expression of the risk genes *ALDH1L1* and *IKZF1*, both of which are glia-related genes, was significantly increased in the presence of microglia when stimulated with IFN-γ (Fig.7C). IFN-γ significantly altered the transcriptional program of cerebral organoids and upon comparing the differentially expressed genes between CO and COiMg, we found that most genes show changes in similar directions. However, a proportion of genes showed contrasting behavior in COiMg when exposed to IFN-γ (Fig.7D). Notably, *IGF-1*, *TREML2*, and *AMBN* genes, found downregulated in CO, showed upregulation or no significant change in expression in COiMg (Fig.7E). TREML2, known as triggering receptor expressed on myeloid cells like 2, is expressed by both myeloid and lymphoid cells and is involved in the innate and adaptive immunity in response to inflammation (Ford & McVicar, 2009). AMBN, also known as Ameloblastin, is expressed by endothelial and immune cells, specifically found enriched in microglia cells. It functions as an extracellular matrix protein, promoting adhesion, tissue differentiation and calcium binding (Sjöstedt et al., 2020; Zhang et al., 2011). Downregulation of *TREML2* and *AMBN* in CO followed by its subsequent upregulation in COiMg in response to IFN-γ may indicate an enhancement of microglial immune functions in response to the inflammatory insult. Furthermore, the presence of microglia appeared to partially, or fully, rescue these gene expression changes. The bulk RNA sequencing also revealed a series of upregulated genes in IFN-γ-treated CO that were not differentially expressed in COiMg (Fig.7F). These genes include those associated with glia cells, such as *APLNR*, *HPSE2* and *BARHL1*, primarily expressed by astrocytes, as well as *OLIG1*, *OLIG2*, *OLIG3*, *NKX2-2* and *SKOR2*, predominantly linked to oligodendrocytes. As previously mentioned, the presence of microglia influences the expression of ciliated mesenchymal cell-related genes, by downregulating *FOXJ1*, *CLDN5* and *TTR*. To investigate whether IFN-γ affects their gene expression and whether this is dependent on the presence of microglia, we examined their levels in CO and COiMg. While *FOXJ1*, *CLDN5* and *TTR* were significantly downregulated in CO, their expression did not change significantly in COiMg when exposed to IFN-γ compared to the untreated samples (Fig.7G). Immunofluorescence imaging confirmed a significant decrease in protein expression in IFN-γ-treated CO, whereas no significant change was observed in COiMg (Fig.7H,I,J and Fig.S11A,B,C).

**Figure 7.**
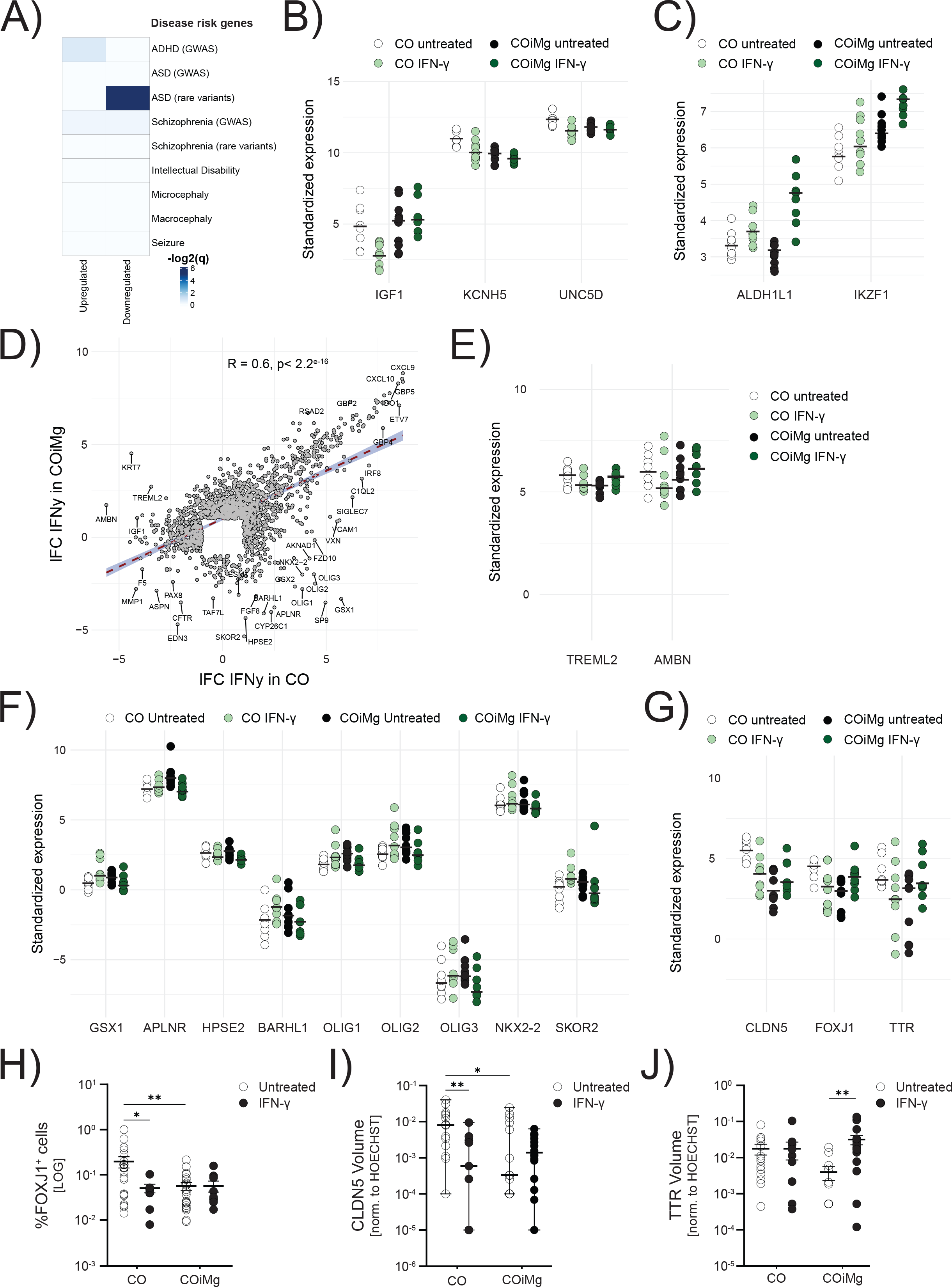
Transcriptional and proteomic changes between CO and COiMg when treated with IFN-γ. (A) Heatmap displaying the -log2(q-values) from GSEA testing overlap between IFN-γ- related DEGs and SFARI databases for psychiatric disorders. Higher -log2(q-values) is equivalent to higher significance of overlap. (B,C) Dotplot of vst-standardized and corrected bulk RNAseq expression values of SFARI ASD rare variants that were also differentially expressed under IFN-γ stimulation in CO and COiMg. (D) Scatter plot displaying the Pearson’s *r* correlation of log-fold changes (filtered for lFC > 1 and lFC < -1) between CO and COiMg treated with IFN-γ. Each dot represents one gene. (E,F) Dotplots of vst-standardized and corrected bulk RNAseq gene expression of selected DEGs between CO and COiMg, when treated with IFN-γ. (G) Dotplots of vst-standardized and corrected bulk RNAseq gene expression of *FOXJ1*, *CLDN5* and *TTR* between CO and COiMg, when treated with IFN-γ. (H) Percentage of FOXJ1^+^ cells was quantified in CO and COiMg. Data are represented as points and error bars as mean ± SEM. N= 2 biological replicates (batches) per line. All three lines pulled. Two-way ANOVA test, followed by Fisher’s LSD post-hoc test, was performed. * P ≤ 0.05, ** P ≤ 0.01. (I) Quantification of CLDN5 surface volume, normalized to HOECHST volume, was performed in CO and COiMg. Data, expressed on a logarithmic scale, are represented as points and error bars as mean ± SEM. N= 2 biological replicates (batches) per line. All three lines pulled. Two-way ANOVA test, followed by Fisher’s LSD post-hoc test, was performed. * P ≤ 0.05, ** P ≤ 0.01. (J) Quantification of TTR surface volume, normalized to HOECHST volume, was performed in CO and COiMg. Data, expressed on a logarithmic scale, are represented as points and error bars as mean ± SEM. N= 2 biological replicates (batches) per line. All three lines pulled. Two-way ANOVA test, followed by Fisher’s LSD post-hoc test, was performed. ** P ≤ 0.01.

## Discussion

Microglia are important for brain development and at regulating the response of the brain to potential harmful insults (Cunningham et al., 2013; Kettenmann et al., 2011). Many central nervous system pathologies (Buonfiglioli & Hambardzumyan, 2021; Hickman et al., 2018), including NDDs (Lukens & Eyo, 2022), have been linked to lack or dysfunction of microglia (Erblich et al., 2011; Oosterhof et al., 2019). iPSC-derived human brain organoids are a valuable tool for modeling the human brain, particularly during development. Most brain organoids predominantly consist of ectodermal lineage-derived cells, neglecting mesodermal lineage-derived cells like microglia and endothelial cells. However, the scientific community has recently focused on integrating brain organoids with these other cell types to elucidate their contribution to brain development and disease (Cakir et al., 2022; Fagerlund et al., 2021; Ormel et al., 2018; Park et al., 2023; Pellegrini et al., 2020; Popova et al., 2021; Schafer et al., 2023).

We generated a model of microglia-containing cerebral organoids (COiMg) that exhibit cells expressing typical microglial markers, such as IBA1, CD45, TREM2, CD11b, CX3CR1 and, at low levels, TMEM119, which are capable of responding to inflammatory stimuli. Morphologically, microglia in COiMg appear amoeboid with few ramifications. The morphology and the microglial marker expression pattern in our organoids is in line with other protocols, suggesting that microglia in the brain organoid environment adopt the microglial phenotype at embryonic stages (Fagerlund et al., 2021; Ormel et al., 2018; Park et al., 2023; Popova et al., 2021). Microglia have been shown to influence neurogenesis, neuronal structure, and activity in the human brain (Cunningham et al., 2013; Thion et al., 2018). We found that the integration of microglia resulted in decreased cell density, decreased neural rosette number and a lower proportion of mature neurons, suggesting that the presence of microglia has an impact on the development of the neuronal compartment, in line with other studies (Fagerlund et al., 2021; Park et al., 2023). Interestingly, we also found that integration of microglia cells in cerebral organoids led to a downregulation of multiciliated mesenchymal cell markers, including CLDN5, TTR and FOXJ1, all involved in endothelial and ependymal cell differentiation and choroid plexus formation. Our results suggest that microglia also influence the development of endothelial and ependymal cells in the organoid, in line with previous research showing a role for microglial cells in brain vasculature development (Hattori, 2023).

We used the CO and COiMg model to investigate the impact of early immune challenges on the developing human brain (Hall et al., 2023). We first screened the response of the organoids to a panel of inflammatory triggers, and found that microglia influence some, but not all, inflammatory responses of cerebral organoids. Next, we investigated the transcriptome response to LPS and IFN-γ. In contrast to the protein analysis, where we found increased release of several cytokines after LPS exposure, we did not find differential expressed genes 48 hours after LPS stimulation. The response of immune cells to LPS is known to be fast, reaching a peak within hours (Das et al., 2016). Our results indicate that this is also the case for organoids, and that the response to LPS was likely already dampened. As we were interested in understanding the longer impact of inflammatory insults on the developing brain, we continued characterizing the response to IFN-γ.

IFN-γ is found at high levels in maternal serum during viral infections or stress (Goines et al., 2011; Ivashkiv, 2018; Jones et al., 2017), as well as in the plasma of neonates who will develop and be diagnosed with ASD (Goines et al., 2011; Heuer et al., 2019; Sasayama et al., 2017; Singh, 1996). We confirmed that IFN-γ can have an effect on the developing brain, using our cerebral organoid model. Notably, we found an overlap between the responses of CO and COiMg, indicating that non-immune cells also respond to this cytokine. IFN-γ has been shown to exhibit a dual role in the brain, exerting both beneficial and detrimental effects (Ottum et al., 2015) depending on the concentration used as well as the specific target cell population. In our cerebral organoid model, IFN-γ appeared to significantly increase the number of nuclei only in COiMg. This finding may potentially align with previous murine studies demonstrating that low doses of IFN-γ induce neurogenesis and oligodendrogenesis via microglial activation (Butovsky et al., 2006). Another explanation of increased cell density may be due to the observed increased microglial number after IFN-γ application, as already seen in previous animal studies (Ta et al., 2019). Furthermore, the percentage of SOX2^+^ cells decreased in CO, in line with previous research indicating a direct effect of IFN-γ on neural precursor populations by inhibiting neurogenesis (Li et al., 2010). Interestingly, this decrease in SOX2^+^ cells was no longer significant in COiMg in response to IFN-γ, further supporting a protective role of microglia in our cerebral organoid model. The reduced number of NEUN^+^ cells observed in both CO and COiMg may suggest a potential impact of IFN-γ on mature neurons, regardless of the presence of microglia.

GO pathway analysis revealed that while the IFN-γ-mediated response is highly similar between CO and COiMg, the intensity of the upregulated biological processes-related genes is increased in cerebral organoids containing microglia. Additionally, the downregulated biological pathways differ between CO and COiMg. Interestingly, we found an upregulation of the ER-pathways in IFN-γ-treated CO but not COiMg. This is in line with previous RNAseq studies showing an upregulation of ER stress related-pathways in maternal immune activation models of human iPSC-derived neurons treated with IFN-γ (Kathuria et al., 2022) and in poly(I:C)-exposed murine fetal brains (Kalish et al., 2021). Moreover, both at the transcriptional and protein levels, microglia appear to modulate the expression of several genes in response to IFN-γ, including those associated with glia and cilia, indicating a regulatory role of microglia in early immune challenges in cerebral organoids.

Finally, exposing cerebral organoids to IFN-γ appeared to be associated with an altered expression of genes linked to ASD risk. In organoids without microglia, IFN-γ seemed to downregulate certain disease risk genes, such as IGF-1, whose expression and signaling are altered in ASD (Riikonen, 2016; Riikonen et al., 2006). Microglia are the major source of IGF-1 and recent animal studies have even proposed a role for microglia-derived IGF- 1 on proper brain development and synaptic functionality (Rusin et al., 2024). In our study the integration of microglia rescued the downregulation, suggesting a protective role of microglia against immune insults.

## Strengths, limitations & concluding remarks

We propose our model of cerebral organoids with integrated microglia to model and study the human developing brain and its exposure to immune challenges. This study has several strengths. First, it provides an opportunity to explore the role of microglia in neurodevelopment and to study the organoid response to inflammatory triggers. Second, unlike other protocols where externally differentiated microglia are incorporated into cerebral organoids (Popova et al., 2021), our approach, which involves integrating microglial progenitors, allows for a more natural microglial differentiation process. Similarly, compared to protocols where microglia develop innately within the organoids (Cakir et al., 2022; Ormel et al., 2018) without control in cell ratio and behavior, our co-culture model provides a more controlled approach for the presence of microglia, as well as the manipulation of cell ratio and microglial cell percentage in cerebral organoids. Lastly, we ensured robust and reproducible data by using multiple iPSC donor lines and performing experiments with multiple batches and replicates.

A limitation observed in our model, which has been seen in several protocols, is the variability between organoids and iPSC lines. We provided a transparent depiction of this variability through a comprehensive representation of all collected data points. Recognizing this variability is crucial for designing future experiments. Addressing will entail larger sample sizes and validating results across multiple iPSC cell lines.

Notably, our model addresses the early stages of brain development when microglia enter and shape the brain. At this stage astrocytes are not yet fully mature. Also, the analysis of a single time point can be a limiting factor of this study. Exploring later time points would not only provide insights into the ongoing development of the organoids and the intricate interplay between microglia and other cell types, including astrocytes, but also allow to study the long-term effects of inflammatory triggers on the brain. It would also enable us to assess whether any observed effects of IFN-γ persist or evolve over time, shedding more light on its potential role in neurodevelopmental disorders. Nevertheless, the data presented here offer a first good picture of transcriptomic and protein changes induced by microglia and IFN-γ.

In conclusion, our work provides a platform to study the brain and microglia during neurodevelopment, and to model early-stage exposure to inflammatory triggers.

## Acknowledgements

The work was supported by the NIH/NIMH grant R21-MH120581. We would like to thank the Microscopy and Advance Bioimaging Core at Mount Sinai, the Dean’s Flow Cytometry CORE at Mount Sinai and the Stem Cell Engineering Core at the Icahn School of Medicine at Mount Sinai for the technical assistance and the opportunity to carry out some of the work. We further thank PhD Gonzalo Pinero and PhD Juan Turati for their help and discussion. All illustrations were created with Adobe Illustrator 2022.

## Declaration of interests

The authors declare no competing interests.

## CRediT authorship contribution statement

**Alice Buonfiglioli**: Writing, Conceptualization, Data curation, Formal Analysis, Investigation, Methodology, Project administration, Visualization, Validation. **Raphael Kübler**: Writing, Investigation, Formal Analysis, Software, Data Curation. **Roy Missall**: Investigation. **Renske De Jong**: Investigation, Formal analysis. **Stephanie Chan**: Investigation, Formal analysis. **Verena Haage**: Investigation, Formal analysis, Resources. **Stefan Wendt**: Investigation, Formal analysis, Supervision. **Ada J. Lin**: Investigation, Formal analysis, Software. **Daniele Mattei**: Formal analysis, Supervision. **Mara Graziani**: Investigation, Formal analysis. **Brooke Latour**: Supervision. **Frederieke Gigase**: Investigation. **Haakon B. Nygaard**: Resources. **Philip L. De Jager**: Supervision, Resources. **Lotje D. De Witte**: Writing, Conceptualization, Data Curation, Funding Acquisition, Project Administration, Supervision, Visualization.

## Data availability

The raw RNA sequencing data will be deposited in NCBI’s Gene Expression Omnibus and will be accessible through GEO Series accession number at the time of publication.

## Material and Methods

### Induced Pluripotent Stem Cell Cultures

The human iPSC line WTC11 (identifier GM25256) was purchased from the Allen Institute, while the MSN (Mount Sinai Normal) -38 and -9 lines were obtained from the Stem Cell Engineering Core Facility at Icahn School of Medicine at Mount Sinai (Schaniel et al., 2021). All lines were authenticated by Short Tandem Repeat (STR) analysis, characterized by immunofluorescence microscopy (Nanog, Oct4, Sox2, SSEA4, TRA1-60) and tested negative for Sendai Virus and mycoplasma. The WTC11 line was generated from a healthy male Asian donor and the MSN38 and MSN9 were generated from a female Asian and a white female, respectively. The WTC11-EGFP (AICS-0036-006) was purchased from the Coriell research institute for medical research and expresses mEGFP under the control of a CAGGS promoter, which has been inserted at the safe harbor locus (AAVS1) using CRISPR/Cas9 technology. All the iPSC lines were cultured in a 6-well plate as monolayer on 0.1 mg/mL Matrigel (cat#: 354277; Corning) in KnockOut™ DMEM (1x) (cat#: 10829018; Gibco) media. The cells were maintained in mTeSR™ Plus media (cat#: 100-0276; StemCell Technologies) and kept at 37°C with 5% CO^2^ with fresh media change every other day. Once 60-70% confluency was reached, the cells were split with mechanical dissociation by using 0.5 mM ultrapure EDTA (cat#: 15575020; Invitrogen) in PBS.

### Generation of Cerebral Organoids (CO)

To differentiate iPSCs into cerebral organoids, the STEMdiff™ Cerebral Organoid Kit (cat#: 08570; StemCell Technologies) was used according to the manufacturer’s manual, with some modifications. On day 0, iPSCs were detached with EDTA and digested in single cells using 0.5 mM STEM Pro Accutase (cat#: A1110501; Gibco). Cells were plated at a confluency of 9,000 cells/well in a Ultra-low attachment 96U-well microplate (cat#: 07-201-680; Thermo Fisher Scientific) in Embryoid Body (EB) Formation Medium (STEMdiff™ Cerebral Organoid Basal Medium 1 + Supplement A), containing 10 µM Y- 27632 (cat#: 72302; StemCell Technologies) and 100 µg/mL of the antibiotic Primocin (cat#: ant-pm-1; InvivoGen). The plate was centrifuged at 100 rcf for 3 min and incubated at 37°C for 48h. On day 2 and 4, 100 µL/well of fresh EB Media was added. The EBs were transferred to an Ultra-low attachment 24-well microplate (cat#: 07-200-602; Thermo Fisher Scientific), 1 EB/well, and 400 µL/well of Induction Medium (STEMdiff™ Cerebral Organoid Basal Medium 1 + Supplement B + 100 µg/mL Primocin) was added. On day 7, the EBs were transferred into an Ultra-low attachment 24-well plate in 400 µL/well of Expansion Medium (STEMdiff™ Cerebral Organoid Basal Medium 2 + Supplement C and D + 100 µg/mL Primocin) supplemented with 2% growth factor reduced Matrigel® (cat#: 354230; Corning). On day 10, 4 EBs/well were transferred to an Ultra-low attachment 6-well plate (cat#: 07-200-601; Thermo Fisher Scientific) and maintained in Maturation Medium (STEMdiff™ Cerebral Organoid Basal Medium 2 + Supplement E + 100 µg/mL Primocin) on an orbital Shaker (65 rpm) at 37°C and 5% CO^2^. The COs were maintained in Maturation Media with fresh media change twice a week until further experiments.

### Generation of Hematopoietic Stem Cells (HPC)

For the generation of HPCs the STEMdiff Hematopoietic Kit (cat#: 05310; StemCell Technologies) was used, according to the manufacturer’s instructions. Briefly, when 70% confluent (day 0), iPSCs were harvested and passaged at a density of 20-40 colonies per well in a 6-well plate coated with 0.1 mg/mL Matrigel (cat#: 354277; Corning). On day 1, Medium A was added to the culture and on day 4 it was switched to Medium B, until complete HPC differentiation on day 10-12. Fully differentiated HPCs, detached from the colonies and cells floating in the medium, were collected for further assays, like microglial differentiation or COiMg generation.

### Generation of Microglia-containing Cerebral Organoids (COiMg)

To generate microglia-containing cerebral organoids (COiMg), HPCs were co-cultured for 7 days with COs (age 25-35 days) with Maturation Medium, supplemented with the cytokines IL-34 (cat#: 1021520; PeproTech), TGF-β (cat#: 100-21; PeproTech), and M-CSF (cat#: 300-25; PeproTech) at the concentration of 100ng/mL, 50 ng/mL, and 25ng/mL, respectively. CO without HPCs nor cytokines, were used as reference control. CO and COiMg were cultured in an Ultra-low attachment 24-wells microplate (cat#: 07- 200-602; Thermo Fisher Scientific), at confluency of 1 organoid/well. 500 µL of fresh media per well was added every other day. After 7 days of co-culture, CO and COiMg were either harvested or used for downstream experiments.

### Generation of Organoid-conditioned Microglia (ocMG)

After 7 days of COiMg culture, we harvested the HPCs that were not integrated into organoids and differentiated these cells into Organoid-conditioned Microglia (ocMG). ocMG were cultured in 0.1% Poly-L-lysine (cat#: P8920; Millipore Sigma)-coated plates using a 50-50 combination of fresh Maturation Medium and COiMg-conditioned Media for an additional 7 days, with media feed every other day.

### Generation of iPSC-derived Microglia (iMG)

iPSC-derived microglia (iMG) were differentiated as previously shown (McQuade & Blurton-Jones, 2022; McQuade et al., 2018). Briefly, differentiated HPCs were collected and transferred in Poly-L-Lysine (PLL)-coated 6 well plate at a confluency of 500k HPC per well. The iMG differentiation takes 25-28 days, during which HPC are cultured in iMG medium consisting of DMEM/F12 (cat#: 11320-033; Gibco), with 2X insulin-transferrin-selenite (ITS) (cat#: 41400045; Thermo Fisher Scientific), 2X B27 (cat#: 17504044; Thermo Fisher Scientific), 0.5X N2 (cat#: 17502048; Thermo Fisher Scientific), 1X Glutamax (cat#: 35050061; Gibco), 1X non-essential amino acids (cat#: 11-140-050; Gibco, 400 mM Monothioglycerol (cat#: M6145; Millipore Sigma), and 5 mg/mL human insulin (cat#: I9278; Millipore Sigma), freshly supplemented with 100 ng/mL IL-34 (cat#: 200-34; PeproTech), 50 ng/mL TGFβ1 (cat#: 100-21; PeproTech), and 25 ng/mL M-CSF (cat#: 300-25; PeproTech).

### Isolation of human post-mortem microglia (pMG)

Microglia from human post-mortem brain tissue was extracted according to Mattei et al. (Mattei et al., 2020) with some modifications for human brain. Briefly, occipital brain tissue was minced in a pre-chilled petri dish in ice-cold Hibernate-A, transferred in a 15 mL Dounce homogenizer and mechanically dissociated by using a loose pestle. The tissue homogenate was then passed through a 100 µm strainer in a 50 mL falcon tube and centrifuged at 450 rcf for 10 minutes. Subsequently, the pellet was resuspended in ice-cold DPBS and afterwards mixed well with 7.2-7.4 pH isotonic Percoll solution (7.3-7.4 Percoll in 10X DPBS in a 9:1 dilution). Finally, overlay with ice-cold DPBS was done slowly to create the cell suspensions and the layered sample was centrifuged at 3000 rcf for 10 minutes. Once the supernatant was aspirated, the pellet was gently washed with DPBS and centrifuged at 400 rcf for 10 minutes. Finally, microglia were extracted from the cell suspension via magnetic-activated cell sorting (MACS) using human anti-CD11b (cat#: 130093634; Miltenyi).

### Stimulation of 2D and 3D cultures

iMG and ocMG, as well as CO and COiMg, were stimulated for 6, 24, and 48 hours with the following ligands: LPS (cat#: L2630; Millipore-Sigma), IFN-α (cat#: 111011; Fisher Scientific), IFN-γ (cat#: 285IF100; Fisher Scientific), IL-6 (cat#: 200-06; Fisher Scientific). All ligands were used at a concentration of 100 ng/mL.

### Multiplex ELISA

To characterize the inflammatory response to stimuli in microglia and organoids, supernatants were collected at 6, 24 and 48 hours after adding microglia and tested using a beads-based multiplex ELISA Immunoassay. We used Luminex xMAP technology for multiplexed quantification of 8 human cytokines, chemokines, and growth factors. The multiplexing analysis was performed using the Luminex™ 200 system (Luminex, Austin, TX, USA) by Eve Technologies Corp. (Calgary, Alberta). Eight markers were simultaneously measured in the samples using Eve Technologies’ Human Cytokine Panel A 8-Plex Custom Assay (MilliporeSigma, Burlington, Massachusetts, USA) according to the manufacturer’s protocol. The 8-plex consisted of IL-6, IL-8, IL-10, CXCL10, MCP-1, MCP-3, CXCL9, and TNF-α. Assay sensitivities of these markers range from 0.14 to 8.61 pg/mL for the 8-plex. Individual analyte sensitivity values are available in the MilliporeSigma MILLIPLEX® MAP protocol (HCYTA-60K).

### Immunofluorescence microscopy

CO and COiMg were washed three times with PBS and fixed in 4% paraformaldehyde (PFA) for 4h at 4°C. Then, the organoids were washed three times with PBS, transferred to a 30% sucrose solution for 24h at 4°C, and subsequently into cryopreservation blocks containing Tissue-Tek® O.C.T.™ (cat#: 4583; Sakura, Torrance, CA, USA), snap-frozen in 2-Methylbutane (cat#: M32631; Sigma-Aldrich), and stored in -80°C. 30 µm-thick cryosections were made using a cryostat (Leica CM1860) and placed on Superfrost® Plus slides (cat#: 48311-703; VWR Scientific), air-dried, and stored at −20°C.

Organoid sections were blocked with Blocking Buffer, constituted of 0.3% Triton X-100 (cat#: 9036-19-05; Sigma-Aldrich), 5% Donkey serum (cat#: D9663; Sigma-Aldrich), 5% Goat Serum (cat#: G9023; Sigma-Aldrich), 1% Bovine Serum Albumin (cat#: A4503; Sigma-Aldrich) in PBS, for 2 h at room temperature. Primary antibodies (Supplementary Table 1) were incubated in blocking buffer overnight at 4°C. Afterwards, the slides were washed three times in PBS, and incubated with secondary antibodies (Supplementary Table 1) for 2h at room temperature. Nuclei were visualized with Hoechst 33342. After three washes in PBS, the slides were embedded in FluorSave Reagent mounting media (cat#: 345789; Sigma-Aldrich), dried and stored in the fridge until subsequent imaging.

### Image acquisition and 3D analysis

Immunofluorescence acquisition was done using the SP8 Confocal Microscope (Leica). Images were acquired with a 20X dry objective by using a z-stack with a 1 µm step size interval and tile scan mode. Protein expression was evaluated by using Imaris v10 (Bitplane). The Surface and Spot tools were used to quantify the area and the volumes of the organoids, and the objects corresponding to the specific cell markers. Colocalization analysis was done by using the “Spot to Spot” colocalization method. The surface details settings were the following: surface detail for all channels was 1 µm. DAPI (seedpoint: 2 µm; threshold: 10; quality: 1.10; voxel: 100), GFP (background subtraction threshold: 10; voxel: 200), SOX2 (seedpoint: 3 µm; threshold: 20; quality: 0.5; voxel: 200), NEUN (seedpoint: 5 µm; threshold: 10; quality: 0.5; voxel: 100).

### Flow Cytometry

CO and COiMg were washed three times in ice-cold DMEM/F12 and then mechanically dissociated. The mechanical dissociation was performed in ice-cold DMEM/F12- containing 40 units/mL DNAse (cat#: LK003172; Worthington) and adapted according to Mattei et al. 2020 protocol (Mattei et al., 2020). Briefly, organoids were mechanically dissociated in a 1 mL Dounce homogenizer with a loose pestle (cat#: 40401; Active Motif) by pipetting up and down approximately 10 times. The cell suspension was then passed through a 100 µm strainer and collected in a 15 mL falcon tube. The Douncer was washed twice with 1 mL DMEM/F12, added to the tube and the cells were centrifuged at 300 g, at 4°C for 5 min. Afterwards, cell counting was performed, and cells were resuspended in ice-cold FACS buffer (PBS supplemented with 2% BSA and 0.5 mM EDTA). The cells were first blocked for 10 minutes with Fc Blocker (cat#: BDB564765; BD Biosciences) and subsequently stained with antibodies (Supplementary Table 1) for 30 minutes at 4°C. The samples were acquired on Cytek Aurora (Cytek Biosciences), and the data analyzed using FlowJo v10.10 (Tree Star).

### Total RNA extraction

2D cultures were harvested and centrifuged at 400 rcf for 6 minutes. The cell pellet was resuspended in Qiazol (cat#:79306; Qiagen). CO and COiMg were washed two times in PBS and dissociated in Qiazol. All cells-diluted in Qiazol, were either stored at -80°C or directly processed for total RNA extraction by using the miRNeasy Micro Kit (cat#: 74004; Qiagen) according to the manufacturer’s manual. The RNA purity and concentration was measured with the NanoDrop™ 2000 (Thermo Fisher Scientific).

### Bulk RNA sequencing

RNA samples were quantified using Qubit 2.0 Fluorometer (Life Technologies) and RNA integrity was checked using Agilent TapeStation 4200 (Agilent Technologies).

The RNA sequencing libraries were prepared using the NEBNext Ultra II RNA Library Prep Kit for Illumina using manufacturer’s instructions (New England Biolabs). Briefly, mRNAs were initially enriched with Oligod(T) beads. Enriched mRNAs were fragmented for 15 minutes at 94°C. First strand and second strand cDNA were subsequently synthesized. cDNA fragments were end repaired and adenylated at 3’ends, and universal adapters were ligated to cDNA fragments, followed by index addition and library enrichment by PCR with limited cycles. The sequencing libraries were validated on the Agilent TapeStation (Agilent Technologies) and quantified by using Qubit 2.0 Fluorometer (ThermoFisher Scientific) as well as by quantitative PCR (KAPA Biosystems).

The sequencing libraries were clustered on five flowcell lanes. After clustering, the flowcell was loaded on the Illumina HiSeq instrument (4000 or equivalent) according to manufacturer’s instructions. The samples were sequenced using a 2x150bp Paired End (PE) configuration. Image analysis and base calling were conducted by the Control software. Raw sequence data (.bcl files) generated from the sequencer were converted into fastq files and de-multiplexed using Illumina’s bcl2fastq 2.17 software. One mismatch was allowed for index sequence identification.

### Single-cell RNA sequencing

Single-cell RNA sequencing experiments were performed as shown in Haage et al. 2024 (Haage et al., 2024). Mechanically dissociated cerebral organoids were filtered through a 35 µm strainer into a low protein binding tube (cat#: 13-864-407; Fisher Scientific). Cell solution was centrifuged at 300g, 4°C for 5 min and washed twice with 1X PBS (cat#: 21- 040-CV; Corning). Next, it was filtered and centrifuged again before being resuspended in 60 µL of PBS containing 0.04% BSA (cat#: 60-00020-10; pluriSelect) for counting using disposable counting chambers (Bulldog Bio). Volumes corresponding to the same cell number from each genetic line (MSN38 and WTC11) were mixed 1:1 to achieve equal cell loading for each sample within one single-cell sequencing experiment. For 10x Genomics, 30,000 cells were targeted for cell loading. The single-cell library preparation was constructed using 10x Chromium Next GEM Single Cell 3’ Reagent Kits v3.1 (Dual Index) according to the manufacturer’s manual. Briefly, a total of ∼30,000 cells were loaded on the 10X genomics chromium controller single-cell instrument. Reverse transcription reagents, barcoded gel beads, and partitioning oil were mixed with the cells for generating single-cell gel beads in emulsions (GEM). After the reverse transcription reaction, the GEMs were broken, and cDNA amplification was performed. The amplified cDNA was then separated by SPRI size selection into cDNA fractions containing mRNA derived cDNA (>300bp) which were further purified by additional rounds of SPRI selection. Sequencing libraries were generated from the mRNA cDNA fractions, analyzed and quantified using TapeStation D5000 screening tapes (Agilent) and Qubit HS DNA quantification kit (Thermo Fisher Scientific).

### Bulk RNAseq Data Analysis

RNAseq reads were aligned along the GRCh38/hg38 reference genome via STAR aligner v2.5 (Dobin et al., 2013) and genes were quantified using featureCounts (Liao et al., 2014). RNAseQC (DeLuca et al., 2012) was used to calculate QC metrics using R (R v4.1.2) and included exonic rate, intergenic rate, mapped reads, rRNA rate, genes detected, and mean per base coverage (Supplementary Table 2). Non-protein-coding genes and genes with a lower base mean than 1 were excluded from analyses. Raw gene counts were vst-normalized to shrink count-inflated gene-gene distances. We used principal component analysis (PCA) to detect outliers. No samples were removed.

We tested a range of experimental variables to determine which of these factors were a major source of variation in our gene expression dataset. We applied PCA, linear regression, and variancePartition (v1.24.0) (Hoffman & Schadt, 2016), which uses a linear mixed model to calculate the percentage of variation in expression due to selected covariates in each gene, to detect potential covariates. We selected factors which explained a significant proportion of the variance but were not correlated with status. The covariates that we selected based on these criteria for subsequent analyses were cell line, 260/280, and 260/230.

Differential expression was calculated with DESeq2 (v1.34.0) (Love et al., 2014). We used a significant threshold of FDR < 0.1 as calculated with the Benjamini-Hochberg procedure. Gene ontology (GO) pathway analyses were performed on differentially expressed genes (DEGs) using the clusterProfiler package (v3.0.4) (Yu et al., 2012). GO molecular function, cellular component, and biological process pathways were included. To further annotate DEGs, gene set enrichment analysis (GSEA) on manually selected gene panels was performed. We used four panels to test for microglia function enrichment: ‘Phagocytosis’ (Sierra et al., 2013), ‘Microglia’ (Patir et al., 2019), ‘Neuroinflammatory I’, and ‘MHC Class II’ (Butovsky & Weiner, 2018). Additionally, we used a list of eight panels to test for genetic risk enrichment in IFN-γ stimulation (Supplementary Table 3): ‘ADHD risk genes (based on GWAS)’ (Demontis et al., 2023), ‘Autism spectrum disorder risk genes (based on GWAS)’ (Grove et al., 2019), ‘Autism spectrum disorder risk genes (rare variants)’ (https://gene.sfari.org/), ‘Intellectual disability risk genes’, ‘Microcephaly risk genes’, ‘Macrocephaly risk genes’, ‘Seizure risk genes’ (https://sysndd.dbmr.unibe.ch/), ‘Schizophrenia risk genes (rare variants)’ (Singh et al., 2022), ’Schizophrenia risk genes (based on GWAS)’ (Trubetskoy et al., 2022). Plots were generated using ComplexHeatmap (v1.10.2) (Gu et al., 2016), ggplot2 (v3.3.5) (Wickham & Wickham, 2016), and RColorBrewer (v1.1.2) (Neuwirth & Brewer, 2014) packages.

### Single-cell RNAseq Data Analysis

Single-cell RNAseq FASTQ files were aligned to the human GRCh38 pre-mRNA reference genome with CellRanger v7.0.1 (https://support.10xgenomics.com/). The filtered gene-cell barcode matrices were used for further analysis with the R package Seurat v4.0. Briefly, cells with more than 20% mitochondria reads, less than 100 genes or more than 5000 genes, were filtered from the downstream analysis. Data was normalized, scaled, and corrected with SCTransform (correcting for number of reads per cell). The first ten principal components were used as input for UMAP and SNN graph construction. A 0.1 resolution was used to calculate clusters with the Louvain algorithm. Clusters were annotated with cell-type identity using the following markers: *AIF1*, *C1QA*, *C1QB*, *CSF1R*, *CX3CR1*, *GPR34*, *ITGAM*, *P2RY12*, *TREM2*, *TMEM119* for microglia; *VIM*, *SOX2*, *PAX6*, *NESTIN*, *Ki67*, *CXCR4*, *SOX1* for neuro-progenitor cells (NPCs); *STMN2*, *MYT1L*, *DCX*, *NRXN1*, *NRXN3*, *MAP2*, *TUBB3*, *TUJ1*, *TBR1*, *TBR2*, *SATB2*, *BCL11B*, *NEUROD1*, *RBFOX3*, *CTIP2*, *NEUROD6*, *NEFL*, *NEFM*, *SLC17A6*, *VAMP2*, *NNAT*, *GAP43*, *GAD2*, *SLC32A1*, *EMX1*, *SCGN*, *PVALB* for cortical neurons; and *COLA1A2*, *COL5A2*, *DCN*, *LUM*, *HES1*, *COL1A6A1*, *DLK1*, *GLI3*, *TUBB4B, AC3*, *FOXJ1*, *ARL13B*, *DNPH1*, *CD63*, *CYR61*, *LAMB1*, *CLDN3*, *CLDN5*, *CLDN1*, *CLDN2*, *FOXJ1*, *TTR*, *CDH1* for ciliated mesenchymal cells. We then calculated differentially expressed genes between culturing condition (CO vs COiMg) using dreamlet v1.1.0, adjusting for experimental batch.

### Quantitative Real-Time PCR (RT-qPCR)

The expression of microglial markers was assessed with quantitative Real-Time Polymerase Chain Reaction (RT-qPCR). 1 µg of total RNA was retro-transcribed into cDNA by using the High-Capacity RNA-to-cDNA Kit (cat#: 01205585; Applied Biosystems) according to the manufacturer’s manual. The expression was analyzed with a QuantStudio 7 Flex (Applied Biosystems) using TaqMan® 2x Universal PCR Master Mix (cat#: 4304437 Applied Biosystems). Absolute and relative RNA expression was determined using the ΔCt and ΔΔCt method, respectively. B-actin was used as a housekeeper gene. The following TaqMan primers were used: ACTB (Hs01060665 g1), AIF1 (Hs00610419_g1), CX3CR1 (Hs00365842_m1), P2RY12 (Hs01881698_s1), TMEM119 (Hs01938722_u1), TREM2 (Hs00219132_m1).

### Rosette Quantification

Rosettes were identified based on nuclei and SOX2 staining. The absolute number of rosettes was manually counted. The average number of rosettes was calculated by dividing the number of rosettes by the surface area of the organoid slice (mm^2^). A representative picture of a rosette morphology is shown in Figure S3B.

### Multi-electrode Array (MEA)

Organoids with and without microglia were plated on a 48-well CytoView MEA plate and recordings were performed on the Maestro Pro MEA recorder from Axion Biosystems. While recording, plates were incubated with 5% CO^2^ and at 37^°^C. The recordings lasted 5 minutes and were performed twice a week, before the media change. At day 51 (post-co-culturing) Tetrodotoxin (TTX) (cat#: 1069; Tocris) at a concentration of 1 µM was used to block any neuronal activity.

Activity from 16 electrodes in each well was recorded simultaneously for five minutes with a sampling frequency of 12.5 kHz. Recordings were subjected to a Butterworth filter of 200-2000 Hz. Spike lists were compiled from the raw recordings, with each spike defined as when the voltage exceeded a threshold of 6 times the standard deviation of the baseline noise of each electrode.

Active electrodes, firing rate, and burst frequency were all calculated from the spike lists using R programming. Active electrodes were defined as any electrode that had a minimum of 5 spikes per minute. The firing rate was calculated as the number of spikes divided by the recording duration in seconds, then normalized to the number of active electrodes in each well. Synchronized bursts were defined as any group of at least 5 spikes having an inter-spike interval of 0.1 seconds or less, for each electrode. Burst frequency was then calculated as the total number of bursts divided by the recording duration in seconds, then normalized to the number of active electrodes in each well.

### Quantification and Statistics

All experiments, from each line, were performed at least three times, unless stated otherwise. All data are represented as mean ± SEM. Statistical analysis was done by using GraphPad Prism (v.10.1; GraphPad Software). Unpaired t-test was used when comparing two groups, while one and two-way ANOVA test was performed to compare multiple groups in case of two variables. Significance was defined at P ≤ 0.05. Except for the RNA sequencing results, only statistically significant P values are shown in the plots. For the RNA sequencing analysis correlation tests were performed in Pearson (nominal) with the basic R stats package (v3.6.2). A *p*-value < 0.05 was considered statistically significant. Principal components were calculated with the prcomp function and simple linear modeling was used to calculate the variance explained by covariates in the principal components using the lm function from the stats package. Differential gene statistical analyses were performed in R (v4.1.2) (CRAN: https://www.r-project.org). We calculated *q*-values using the Benjamini-Hochberg procedure to adjust *p*-values for multiple testing. The Benjamini-Hochberg procedure was also applied to correct for multiple testing in the statistical analyses of the gene panels. GSEA was performed by calculating odds ratios with Fisher’s exact test. We considered a *q*-value of < 0.1 as statistically significant (referred to as FDR < 0.1). All p-values, listed in Supplementary Table 4, were generated from the following comparisons: “CO *versus* COiMg in untreated”, “CO *versus* COiMg in IFN-γ”, “untreated *versus* IFN-γ in CO”, and “untreated *versus* IFN-γ in COiMg”.

---

Supplementary Figure 1. FACS analysis of CO and COiMg for microglial markers

(A) Schematic gating strategy for cell surface marker analysis via FACS. Gating for cells, singlets, and alive cell populations.

(B) Histogram plots depicting the macrophage markers CD14 and CD206.

Supplementary Figure 2. Microglial gene expression and GO analysis in CO and COiMg.

(A) Relative expression of the microglial gene *AIF1* was analyzed via qPCR in COiMg compared to CO in MSN38 and WTC11 iPSC lines. Expression was calculated as ΔΔCT. Data are shown as bars and points, error bars as mean ± SEM. N=4. Unpaired t-test was performed. * P ≤ 0.05, ** P ≤ 0.01.

(B) Absolute expression of the microglial markers *AIF1*, *CX3CR1*, *TREM2*, *P2YR12*, *TMEM119* was analyzed via qPCR in iMG, ocMG, pMG and COiMg from WTC11 and MSN38 iPSC donor lines. Expression was calculated as 2^-ΔCT^.Data are shown as points and error bars as mean ± SEM.

(C,D) Dotplot of Gene Ontology analysis of the up- (C) and down-regulated (D) genes when comparing CO vs COiMg. The size of the dot represents the number of hits in the respective GO term and the color represents the adjusted p-value size.

(E) Heatmap of the Fisher’s exact test -log2(q-values) of the overlap between differentially expressed genes and immune-related pathways relevant to microglia. Larger -log2(q- values) represents higher overlap between the tested gene sets.

Supplementary Figure 3. Structural and functional analysis of CO and COiMg.

(A) Quantification of number of nuclei per mm^2^ in CO and COiMg, derived from MSN38, WTC11 and MSN9 iPSC donor lines. Data are represented as points and error bars as mean ± SEM. N= 2 biological replicates (batches) per line. Unpaired t-test with Welch’s correction was performed.

(B) Schematic representation for the quantification of the rosettes.

(C) Quantification of number of rosettes per organoid area was performed in CO and COiMg, derived from MSN38, WTC11 and MSN9 iPSC donor lines. Data are represented as points and error bars as mean ± SEM. N= 2 biological replicates (batches) per line.

Two-way ANOVA followed by Sidak’s post-hoc test was performed. * P ≤ 0.05.

(D,E) Percentage of SOX2^+^ (D) and NeuN^+^ (E) cells was calculated in CO and COiMg derived from MSN38, WTC11 and MSN9 iPSC donor lines. Data are represented as points and error bars as mean ± SEM. N= 2 biological replicates (batches) per line.f

(F) Dotplot showing the percentage and averaged expression of TNFR1 and TNFR2 in cortical neurons in CO and COiMg.

(G) Raw traces of electrical recordings of cerebral organoids without (left) and with (right) microglia. Resulting patterns of electrical activity are shown below, together with single cell spiking (black) as well as synchronized burst firing (blue).

(H) The overall firing rate (normalized to active electrodes) expressed in Hz was analyzed in CO and COiMg. Data are depicted as bars and error bars as mean ± SEM. N= 1 biological replicate (batch) with 4 CO and 11 COiMg.

(I) Burst frequency was analyzed upon TTX (1µM) application to block neuronal activity during MEA recordings. N= 10 CO (white) and 18 COiMg (black). Wilcoxon test was performed. ** P ≤ 0.01.

Supplementary Figure 4. Cilia-associated gene and protein expression of CLDN5, FOXJ1 and TTR decrease in the presence of microglia in cerebral organoids.

(A) Heatmap of expression Z-statistics displaying whether selected genes (*FOXJ1*, *TTR*, and *CLDN5*) are (de-)enriched across cell-types. Statistical significance, set below 0.05 adjusted p-value, is indicated with a *.

(B) Heatmap of fraction of cells expressing *FOXJ1*, *TTR*, and *CLDN5*. Darker blue indicates a higher fraction of cells expressing the respective genes.

(C) Percentage of FOXJ1^+^ cells was quantified in CO and COiMg, derived from MSN38, WTC11 and MSN9 iPSC donor lines. Data are represented as points and error bars as mean ± SEM. N= 2 biological replicates (batches). Mann Whitney t-test was performed. * P ≤ 0.05.

(D) Quantification of CLDN5 surface volume, normalized to HOECHST volume, was performed in CO and COiMg, derived from MSN38, WTC11 and MSN9 iPSC donor lines. Data are represented as points and error bars as mean ± SEM. N= 2 biological replicates (batches). Mann Whitney t-test was performed. ** P ≤ 0.01.

(E) Quantification of TTR surface volume, normalized to HOECHST volume, was performed in CO and COiMg, derived from MSN38, WTC11 and MSN9 iPSC donor lines. Data are represented as points and error bars as mean ± SEM. N= 2 biological replicates (batches).

(F) Representative image of CO stained with CLDN5 and TTR. HOECHST was used for nuclei staining. 10X Magnification. Scale bar, 100µm.

Supplementary Figure 5. Characterization of the inflammatory landscape of cerebral organoids treated with LPS, IFN-α and IFN-γ.

(A) COiMg, from MSN38 iPSC line, were treated for 24 and 48 h with LPS, IFN-α and IFN-γ (all at 100ng/mL). Subsequently, supernatants were collected and tested for several inflammatory molecules. Untreated COiMg were used as negative control. Release was expressed as pg/mL on a logarithmic scale. Data are shown as bars and points and error bars as mean ± SEM. N=3 biological replicates (batches).

(B,C,D) CO and COiMg, derived from MSN38, WTC11 and MSN9 iPSC donor lines, were stimulated for 48 h with LPS (B), IFN-γ (C) and IFN-α (D), supernatants were collected and cytokine and chemokine release was analyzed via Multiplex ELISA. Data, expressed as fold change release compared to untreated conditions, are shown as bars and error bars as mean ± SEM. N= 3 biological replicates (batches) per line.

Supplementary Figure 6. RNAseq analysis of CO and COiMg treated with IFN-γ.

(A) PCA plot of the first two PCs displaying CO sample distances in untreated (white) and IFN-γ (green) from bulk RNAseq. N= 3 per stimulants per iPSC donor line. PCs were calculated based on the top 500 variable genes.

(B) Dotplot of vst-standardized and corrected expression of the four most upregulated genes in IFN-γ-treated CO.

(C,D) Heatmap of row-scaled vst-standardized and corrected expression values of the top 40 differentially expressed genes between untreated and IFN-γ-treated CO (A) and COiMg (B) samples. Larger Z-score is equivalent to higher gene expression.

Supplementary Figure 7. IFN-γ-mediated signaling pathways are upregulated in CO and COiMg treated with IFN-γ.

(A) Representative image of CO untreated (upper) or treated with IFN-γ (lower) stained with IDO1. Microglia are labelled with GFP. HOECHST was used for nuclei staining. 20X Magnification. Scale bar, 50 µm.

(B) Percentage of IDO1 expression in CO and COiMg, derived from MSN38, WTC11 and MSN9 iPSC lines. Data are represented as points and error bars as mean ± SEM. N= 2 biological replicates (batches).

(C) Percentage of IDO1^+^ microglia was calculated in COiMg, derived from MSN38, WTC11 and MSN9 iPSC lines. Data are represented as points and error bars as mean ± SEM. N= 2 biological replicates (batches).

(D) Representative image of CO untreated (upper) or treated with IFN-γ (lower) stained with IDO1. Microglia are labelled with GFP. HOECHST was used for nuclei staining. 20X Magnification. Scale bar, 50 µm.

(E) Dotplots showing the percentage and averaged expression of IFNGR1 and IFNGR2 in Microglia, Ciliated Mesenchymal cells, Cortical Neurons and NPCs in both CO and COiMg.

Supplementary Figure 8. Cellular and Structural analysis of CO and COiMg treated with IFN-γ.

(A) Quantification of number of nuclei per mm^2^ was performed in untreated and IFN-γ- treated CO and COiMg, derived from MSN38, WTC11 and MSN9 donor lines. Data are represented as points and error bars as mean ± SEM. N= 2 biological replicates (batches) per line.

(B,C) Quantification of number of rosettes per organoid area was performed in untreated and IFN-γ-treated CO and COiMg, derived from MSN38, WTC11 and MSN9 donor lines, pulled (B) or single (C). Data are represented as points and error bars as mean ± SEM.

N= 2 biological replicates (batches) per line. Two-way ANOVA test, followed by Fisher’s LSD post-hoc test, was performed. * P ≤ 0.05.

(D) Percentage of microglia was calculated in untreated and IFN-γ-treated COiMg, derived from MSN38, WTC11 and MSN9 donor lines. Data are shown as points with mean ± SEM. N= 2 biological replicates (batches) per line. Mann Whitney t-test was performed. * P ≤ 0.05.

(E) Percentage of SOX2^+^ cells was calculated in untreated and IFN-γ-treated CO and COiMg, derived from MSN38, WTC11 and MSN9 donor lines. Data are represented as points and error bars as mean ± SEM. N= 2 biological replicates (batches) per line; all donor lines are pulled. Two-way ANOVA test, followed by Fisher’s LSD post-hoc test, was performed. * P ≤ 0.05; ** P ≤ 0.01.

(F) Percentage of NeuN^+^ cells was calculated in untreated and IFN-γ-treated CO and COiMg, derived from MSN38, WTC11 and MSN9 donor lines. Data are represented as points and error bars as mean ± SEM. N= 2 biological replicates (batches) per line; all donor lines are pulled. Two-way ANOVA test, followed by Fisher’s LSD post-hoc test, was performed. ** P ≤ 0.01.

Supplementary Figure 9. Differences in immune-related pathways between CO and COiMg treated with IFN-γ.

(A,B) Heatmap showing the Fisher’s exact test -log2(q-values) of the overlap between differentially expressed genes of untreated *versus* IFN-γ-treated samples and immune- related pathways relevant to microglia in CO (A) and COiMg (B). Larger -log2(q-values) is equivalent to higher overlap.

Supplementary Figure 10. GO analysis in CO and COiMg treated with IFN-γ.

(A,B) Dotplot of GO analysis of the upregulated genes when comparing untreated and IFN-γ-treated CO (A) and COiMg (B). The size of the dot represents the number of hits in the respective GO term and color represents adjusted p-value size.

(C) -log(q) values for selected biological process pathways from the GO analysis of the up-regulated genes in CO (upper) and COiMg (lower).

(D,E) Dotplot of GO analysis of the downregulated genes when comparing untreated and IFN-γ-treated CO (D) and COiMg (E). The size of the dot represents the number of hits in the respective GO term and color represents adjusted p-value size.

(F) -log(q) values for selected biological process pathways from the GO analysis of the downregulated genes in CO (upper) and COiMg (lower).

Supplementary Figure 11. FOXJ1, CLDN5 and TTR expression analysis in CO and COiMg treated with IFN-γ.

(A) Percentage of FOXJ1^+^ cells was quantified in untreated and IFN-γ-treated CO and COiMg, derived from MSN38, WTC11 and MSN9 iPSC donor lines. Data, expressed on a logarithmic scale, are represented as points and error bars as mean ± SEM. N= 2 biological replicates (batches). Two-way ANOVA test, followed by Fisher’s LSD post-hoc test, was performed. * P ≤ 0.05.

(B) Quantification of CLDN5 surface volume, normalized to HOECHST volume, was performed in untreated and IFN-γ-treated CO and COiMg, derived from MSN38, WTC11 and MSN9 iPSC donor lines. Data, expressed on a logarithmic scale, are represented as points and error bars as mean ± SEM. N= 2 biological replicates (batches). Two-way ANOVA test, followed by Fisher’s LSD post-hoc test, was performed. * P ≤ 0.05.

(C) Quantification of TTR surface volume, normalized to HOECHST volume, was performed in untreated and IFN-γ-treated CO and COiMg, derived from MSN38, WTC11 and MSN9 iPSC donor lines. Data, expressed on a logarithmic scale, are represented as points and error bars as mean ± SEM. N= 2 biological replicates (batches).

Supplementary Table 1. Antibody list used for immunofluorescence and FACS.

Supplementary Table 2. Bulk RNAseq QC metrics.

Supplementary Table 3. Annotation gene list.

Supplementary Table 4. DESeq2 DEA results.

Supplementary Table 5. Cilia-associated DEG list.

## Notes

### Competing Interest Statement

The authors have declared no competing interest.

### Summary of Updates

The Acknowledgments and the affiliations have been updated. Abstract updated.

