## Supplementary Figures for "A microglia-containing cerebral organoid model to study early life immune challenges"

### Figure S1

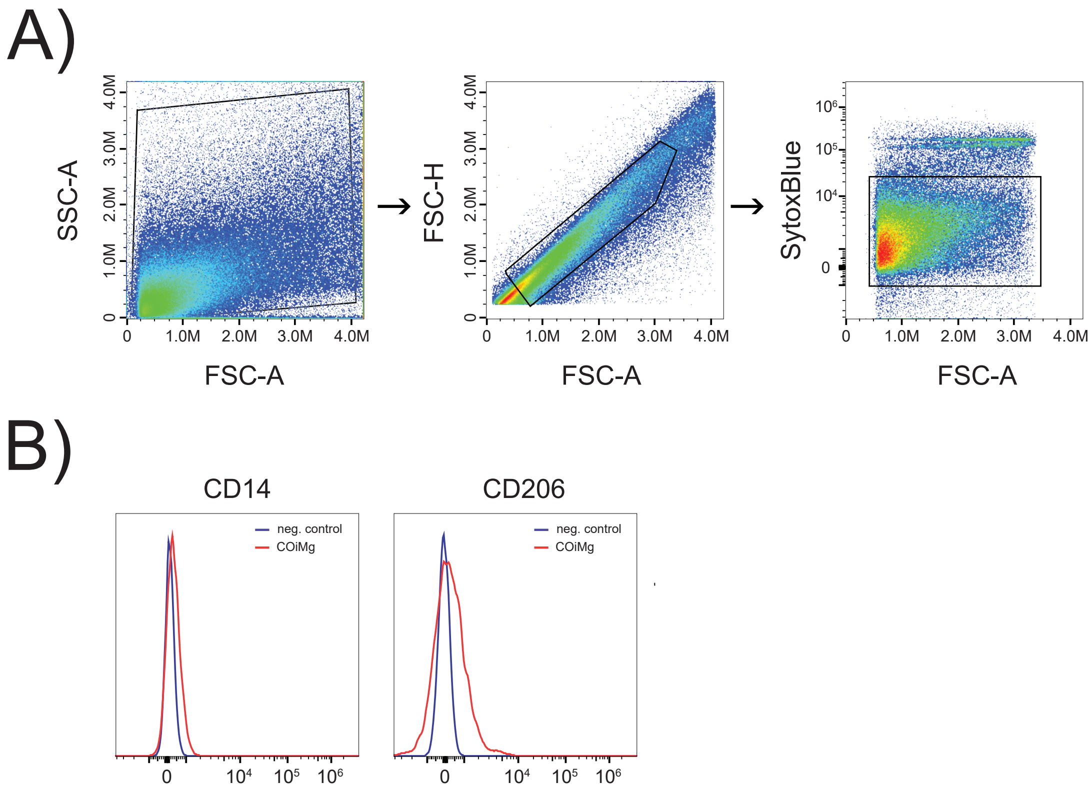

A)
SSC-A
FSC-A
FSC-H
FSC-A
SytoxBlue
FSC-A
B)
CD14
neg. control
COiMg
CD206
neg. control
COiMg

### Figure S2

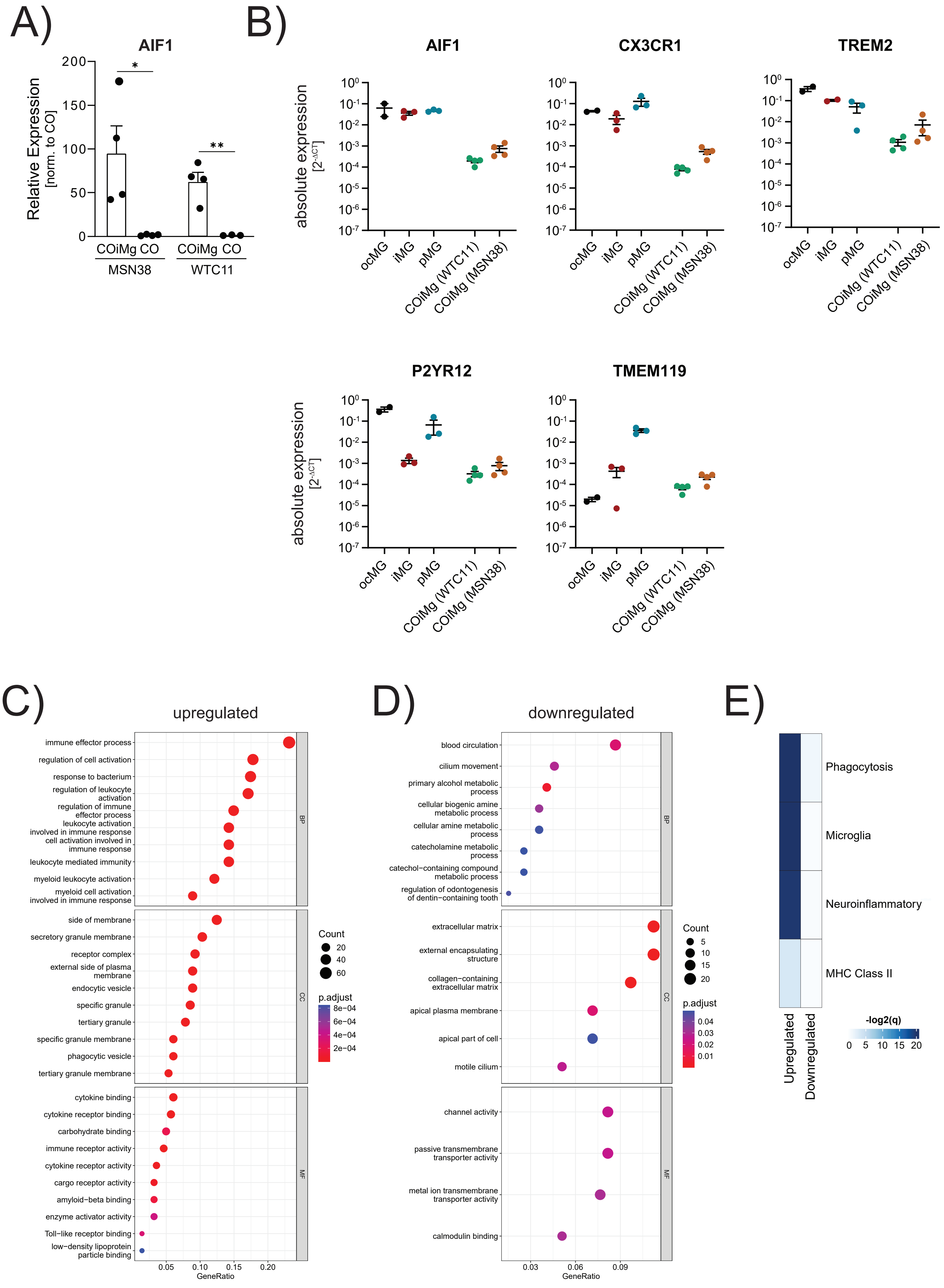

A)
AIF1
Relative Expression
[norm. to CO]
COiMg CO
MSN38
WTC11
B)
AIF1
CX3CR1
TREM2
P2YR12
TMEM119
absolute expression
[2-ΔCT]
ocMG
iMG
pMG
COiMg (WTC11)
COiMg (MSN38)
C)
upregulated
immune effector process
regulation of cell activation
response to bacterium
regulation of leukocyte activation
regulation of immune effector process
leukocyte activation involved in immune response
cell activation involved in immune response
leukocyte mediated immunity
myeloid leukocyte activation
myeloid cell activation involved in immune response
side of membrane
secretory granule membrane
receptor complex
external side of plasma membrane
endocytic vesicle
specific granule
tertiary granule
specific granule membrane
phagocytic vesicle
tertiary granule membrane
cytokine binding
cytokine receptor binding
carbohydrate binding
immune receptor activity
cytokine receptor activity
cargo receptor activity
amyloid−beta binding
enzyme activator activity
Toll−like receptor binding
low−density lipoprotein particle binding
BP
CC
MF
Count
p.adjust
GeneRatio
D)
downregulated
blood circulation
cilium movement
primary alcohol metabolic process
cellular biogenic amine metabolic process
cellular amine metabolic process
catecholamine metabolic process
catechol−containing compound metabolic process
regulation of odontogenesis of dentin−containing tooth
extracellular matrix
external encapsulating structure
collagen−containing extracellular matrix
apical plasma membrane
apical part of cell
motile cilium
channel activity
passive transmembrane transporter activity
metal ion transmembrane transporter activity
calmodulin binding
E)
Phagocytosis
Microglia
Neuroinflammatory
MHC Class II
Upregulated
Downregulated
-log2(q)

### Figure S3

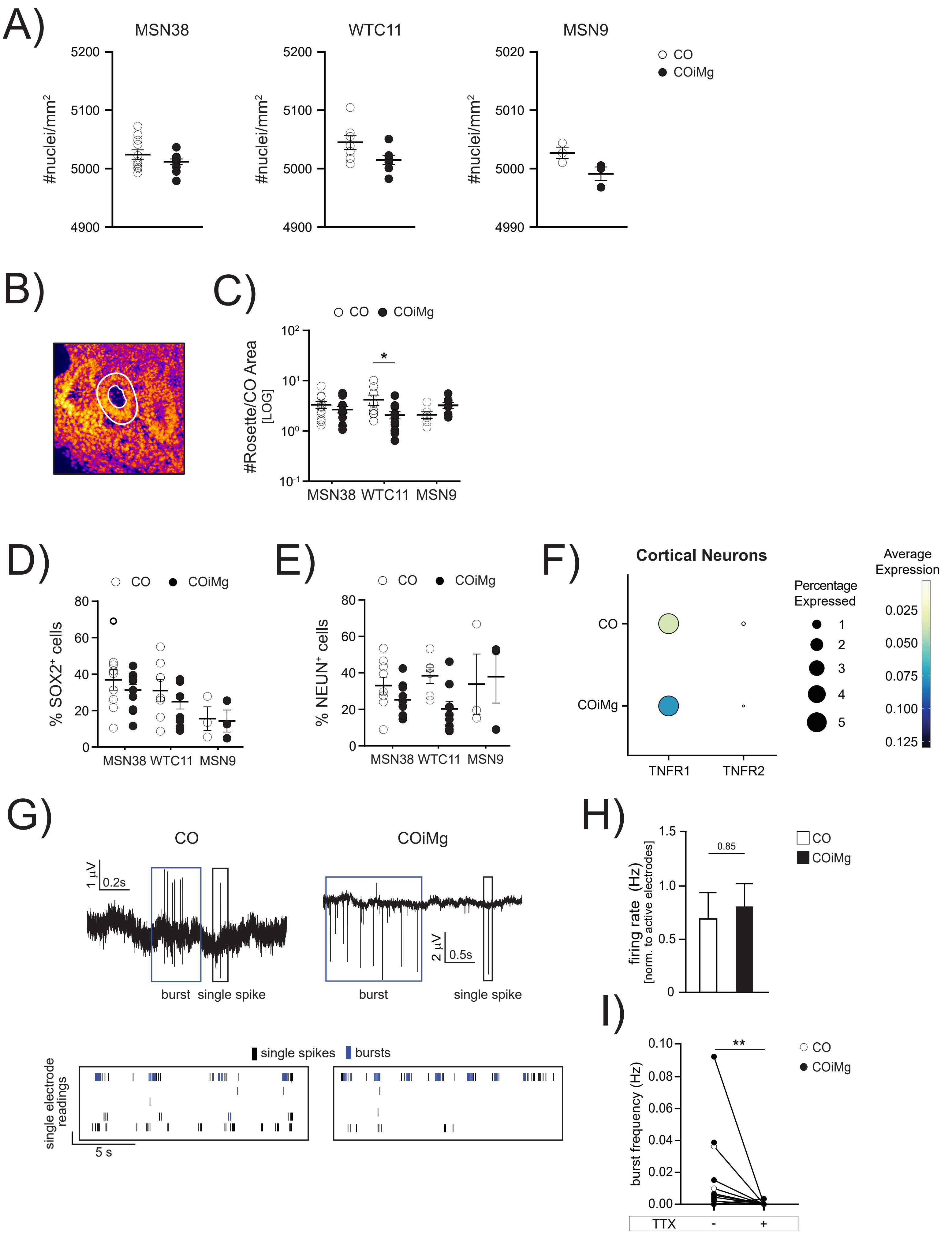

A)
MSN38
WTC11
MSN9
#nuclei/mm²
5200
5100
5000
4900
5020
5010
4990
CO
COiMg
B)
C)
#Rosette/CO Area [LOG]
MSN38 WTC11 MSN9
*
D)
% SOX2⁺ cells
E)
% NEUN⁺ cells
F)
Cortical Neurons
TNFR1
TNFR2
Percentage Expressed
Average Expression
G)
burst
single spike
single spikes
bursts
single electrode readings
5 s
H)
firing rate (Hz)
[norm. to active electrodes]
0.85
I)
burst frequency (Hz)
**
TTX

### Figure S4

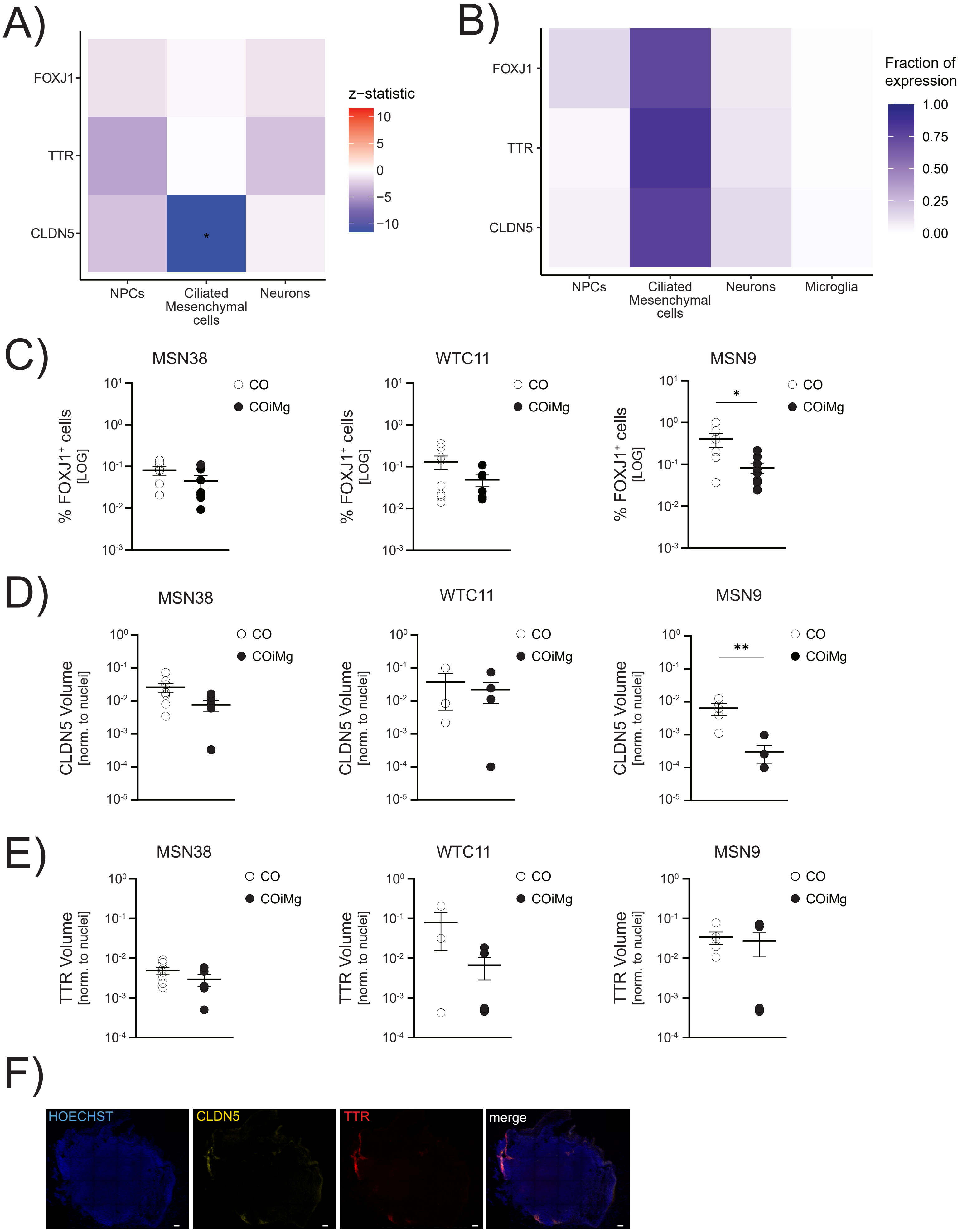
A)
FOXJ1
TTR
CLDN5
NPCs
Ciliated Mesenchymal cells
Neurons
z−statistic
10
5
0
−5
−10
B)
FOXJ1
TTR
CLDN5
NPCs
Ciliated Mesenchymal cells
Neurons
Microglia
Fraction of expression
1.00
0.75
0.50
0.25
0.00
C)
MSN38
WTC11
MSN9
% FOXJ1+ cells [LOG]
CO
COiMg
D)
MSN38
WTC11
MSN9
CLDN5 Volume [norm. to nuclei]
CO
COiMg
E)
MSN38
WTC11
MSN9
TTR Volume [norm. to nuclei]
CO
COiMg
F)
HOECHST
CLDN5
TTR
merge

### Figure S5

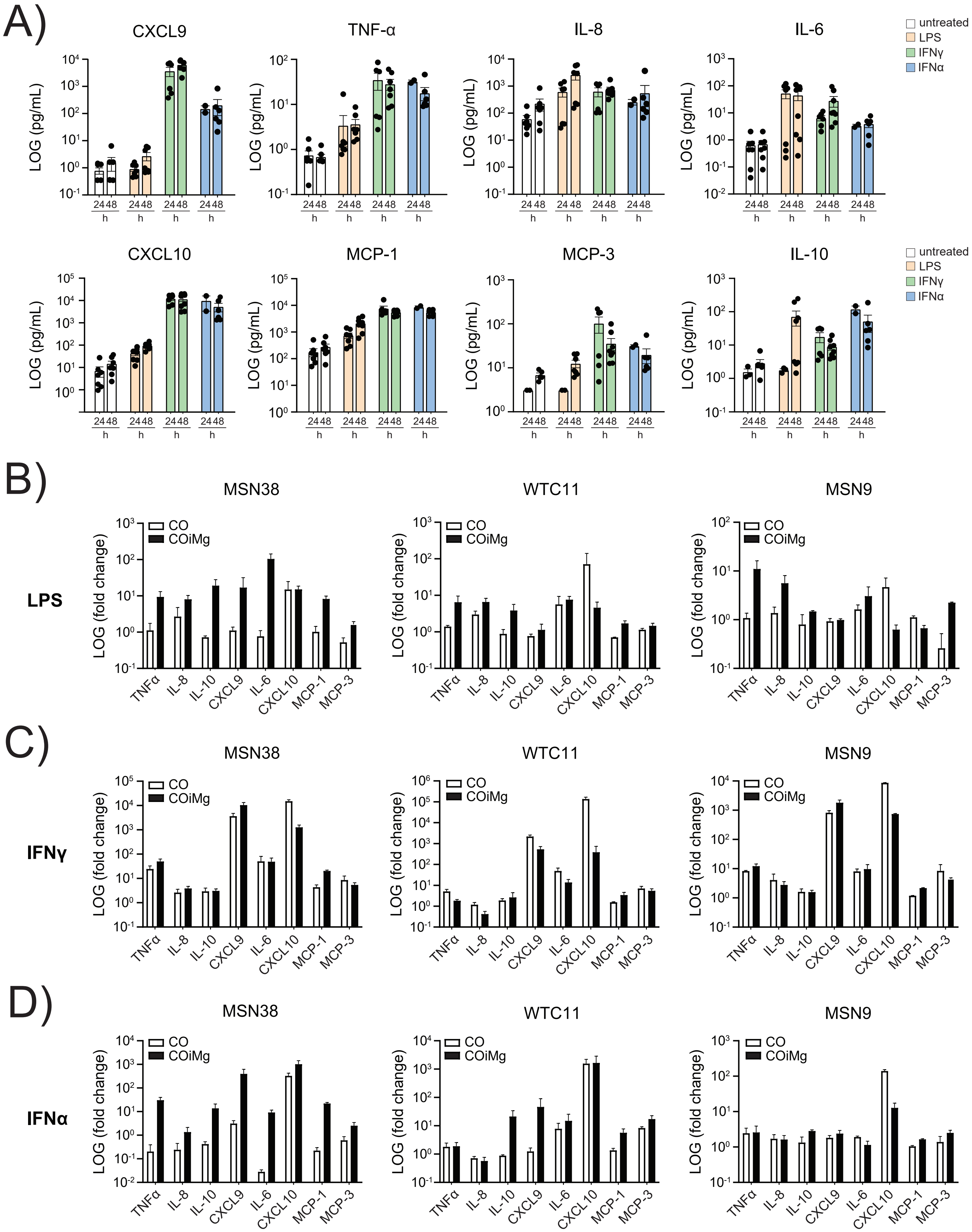
A)
CXCL9
TNF-α
IL-8
IL-6
CXCL10
MCP-1
MCP-3
IL-10
LOG (pg/mL)
24 48 h
untreated
LPS
IFNγ
IFNα
B)
C)
D)
MSN38
WTC11
MSN9
LPS
IFNγ
IFNα
LOG (fold change)
CO
COiMg
TNFα
IL-8
IL-10
CXCL9
IL-6
CXCL10
MCP-1
MCP-3

### Figure S6

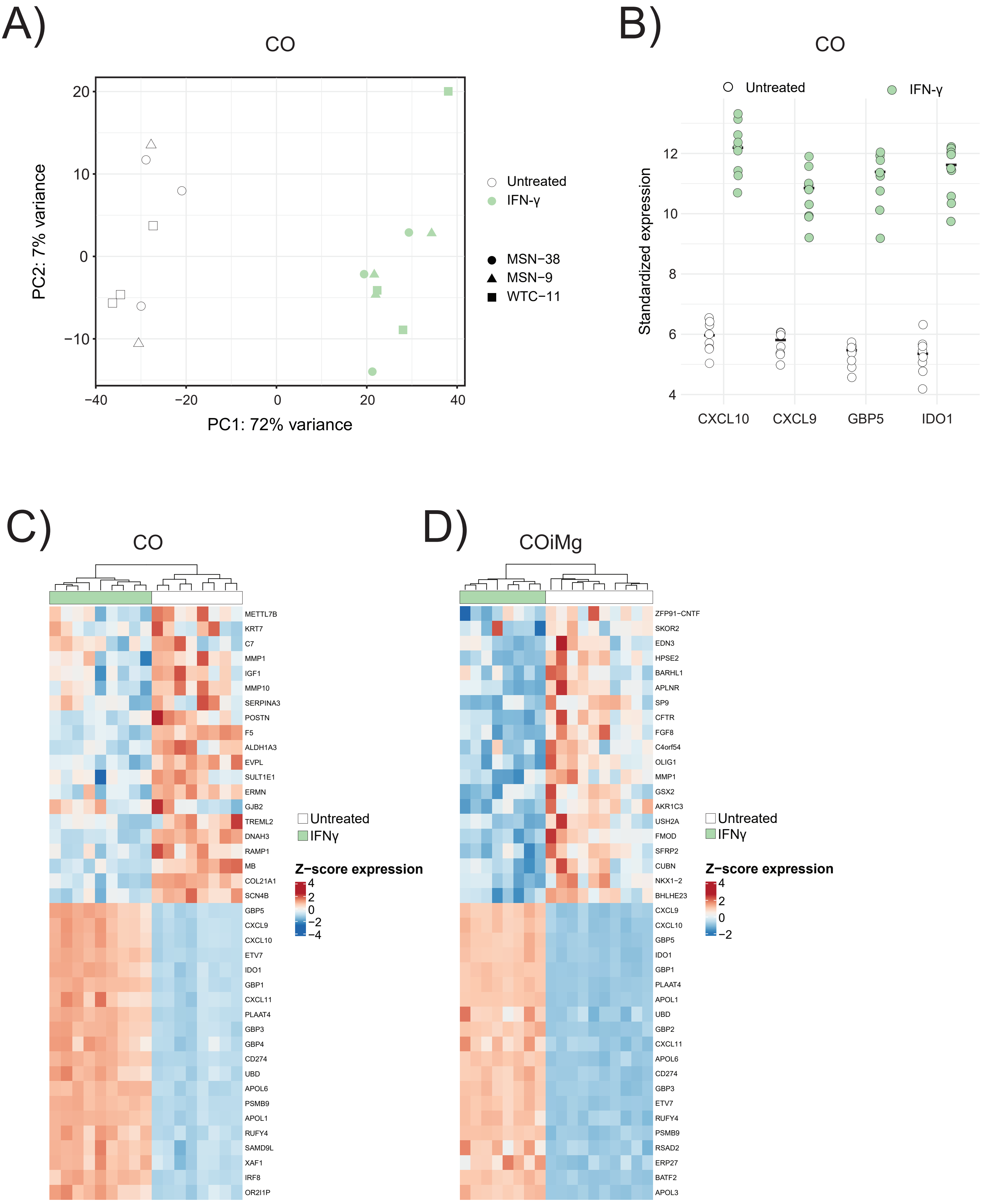

A)
CO
PC1: 72% variance
PC2: 7% variance
Untreated
IFN-γ
MSN−38
MSN−9
WTC−11
B)
CO
Untreated
IFN-γ
Standardized expression
CXCL10
CXCL9
GBP5
IDO1
C)
CO
METTL7B
KRT7
C7
MMP1
IGF1
MMP10
SERPINA3
POSTN
F5
ALDH1A3
EVPL
SULT1E1
ERMN
GJB2
TREML2
DNAH3
RAMP1
MB
COL21A1
SCN4B
GBP5
CXCL9
CXCL10
ETV7
IDO1
GBP1
CXCL11
PLAAT4
GBP3
GBP4
CD274
UBD
APOL6
PSMB9
APOL1
RUFY4
SAMD9L
XAF1
IRF8
OR2I1P
Untreated
IFNγ
Z−score expression
D)
COiMg
ZFP91−CNTF
SKOR2
EDN3
HPSE2
BARHL1
APLNR
SP9
CFTR
FGF8
C4orf54
OLIG1
MMP1
GSX2
AKR1C3
USH2A
FMOD
SFRP2
CUBN
NKX1−2
BHLHE23
CXCL9
CXCL10
GBP5
IDO1
GBP1
PLAAT4
APOL1
UBD
GBP2
CXCL11
APOL6
CD274
GBP3
ETV7
RUFY4
PSMB9
RSAD2
ERP27
BATF2
APOL3
Untreated
IFNγ
Z−score expression

### Figure S7

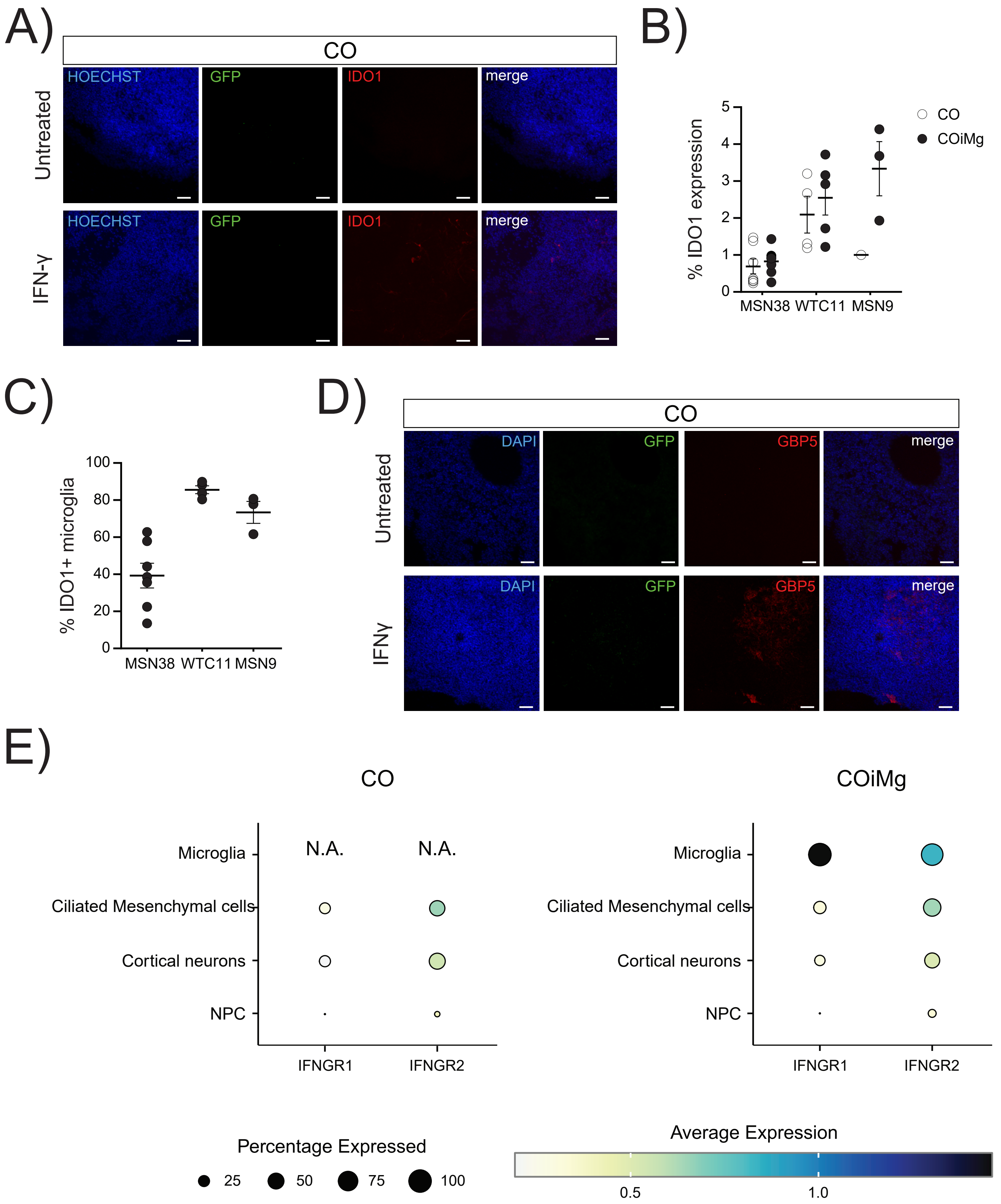

A)
CO
HOECHST
GFP
IDO1
merge
Untreated
IFN-γ
B)
% IDO1 expression
CO
COiMg
MSN38
WTC11
MSN9
C)
% IDO1+ microglia
MSN38
WTC11
MSN9
D)
CO
DAPI
GFP
GBP5
merge
Untreated
IFNγ
E)
CO
COiMg
Microglia
Ciliated Mesenchymal cells
Cortical neurons
NPC
N.A.
N.A.
IFNGR1
IFNGR2
Percentage Expressed
25
50
75
100
Average Expression
0.5
1.0

### Figure S8

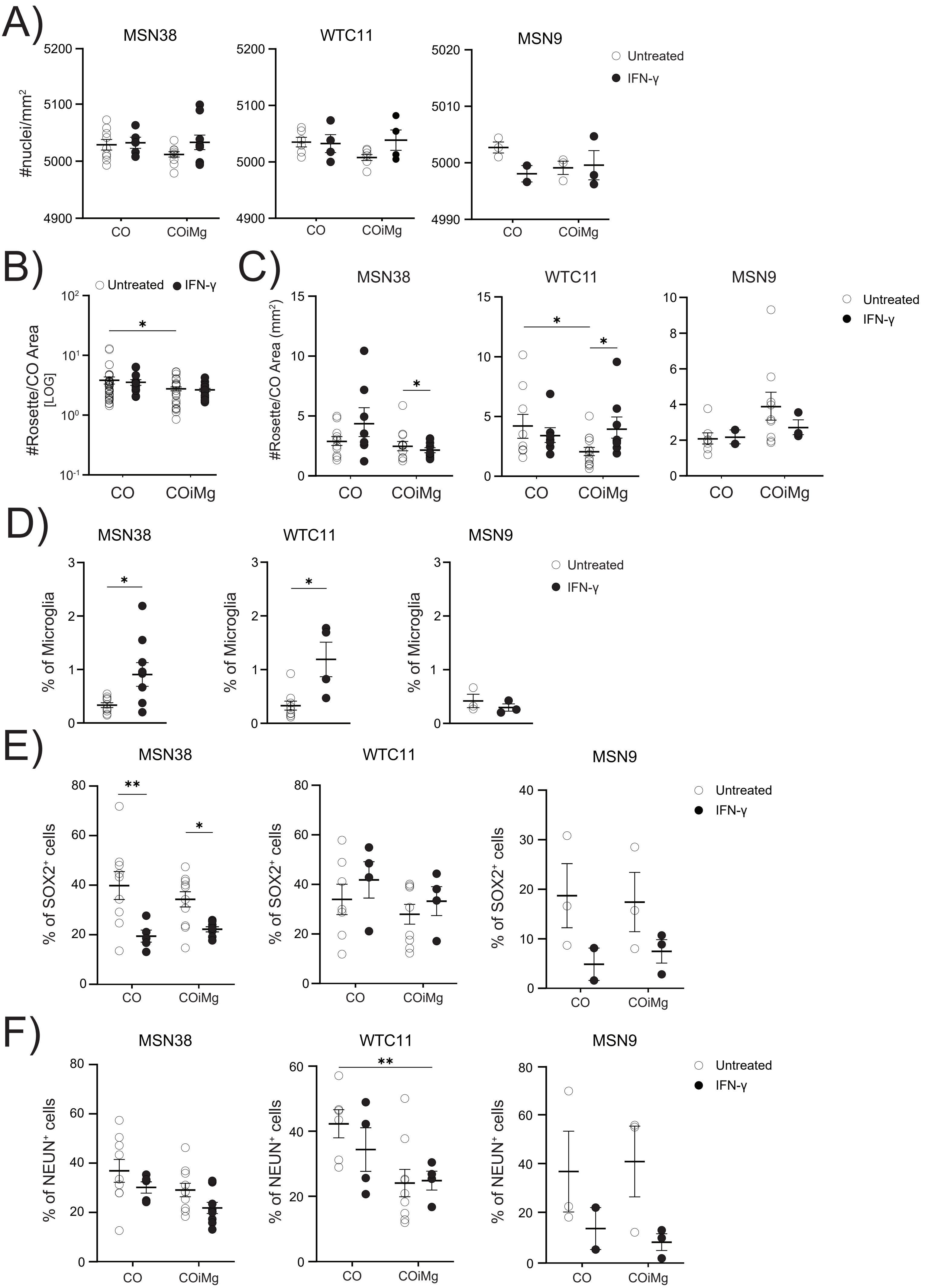
A)
MSN38
WTC11
MSN9
Untreated
IFN-γ
#nuclei/mm²
CO
COiMg
B)
C)
#Rosette/CO Area [LOG]
#Rosette/CO Area (mm²)
D)
% of Microglia
E)
% of SOX2⁺ cells
F)
% of NEUN⁺ cells

### Figure S9

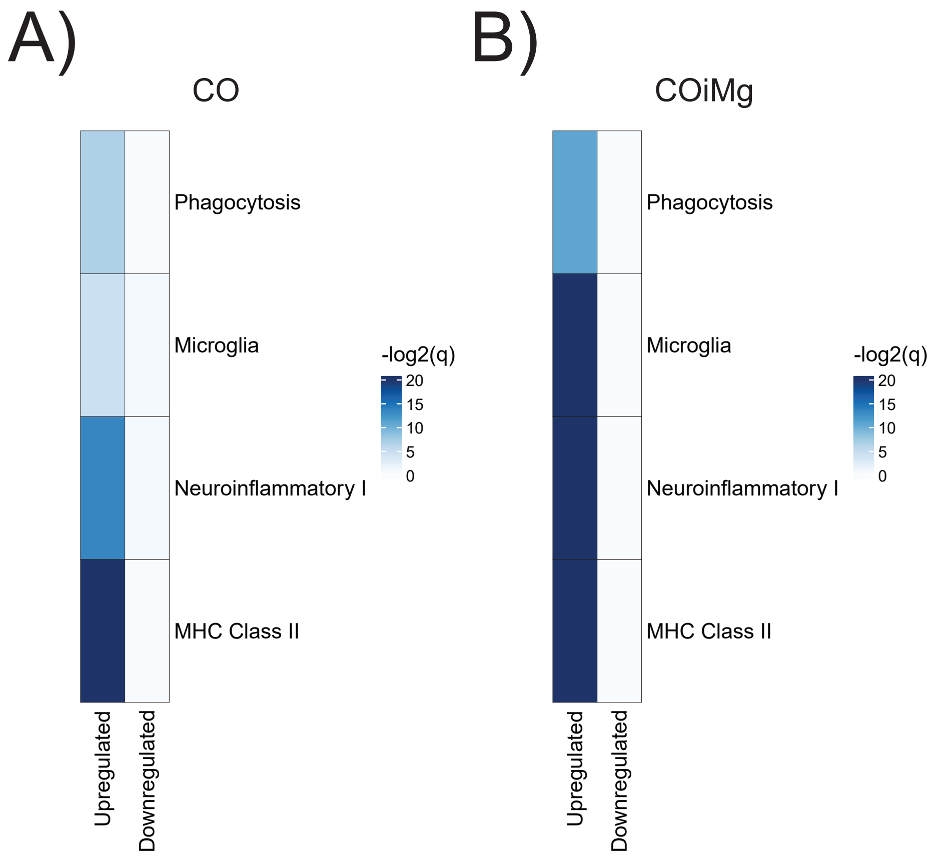
A)
CO
Phagocytosis
Microglia
Neuroinflammatory I
MHC Class II
-log2(q)
20
15
10
5
0
Upregulated
Downregulated
B)
COiMg
Phagocytosis
Microglia
Neuroinflammatory I
MHC Class II
-log2(q)
20
15
10
5
0
Upregulated
Downregulated

### Figure S10

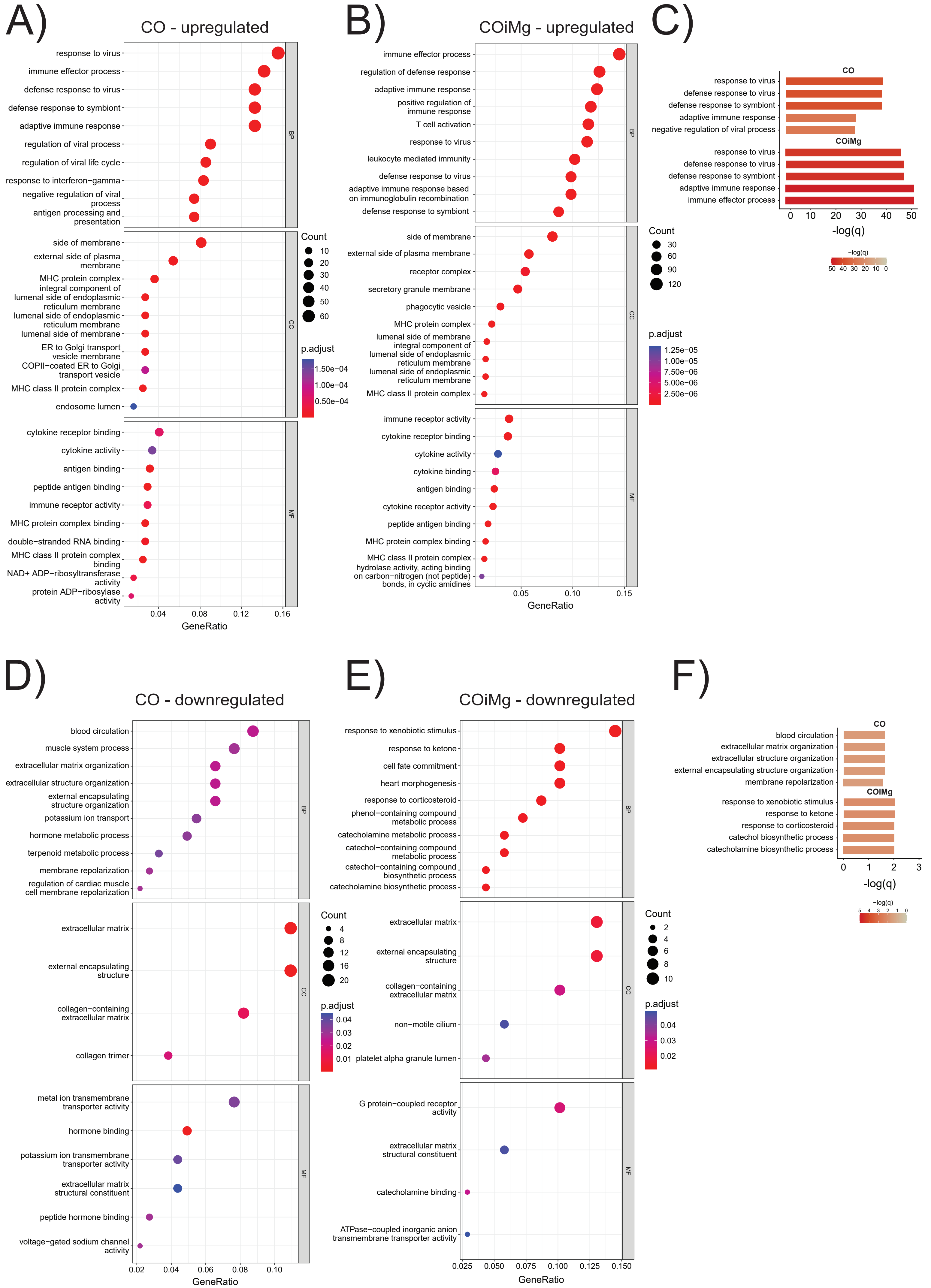
A)
CO - upregulated
response to virus
immune effector process
defense response to virus
defense response to symbiont
adaptive immune response
regulation of viral process
regulation of viral life cycle
response to interferon−gamma
negative regulation of viral process
antigen processing and presentation
side of membrane
external side of plasma membrane
MHC protein complex
integral component of lumenal side of endoplasmic reticulum membrane
lumenal side of endoplasmic reticulum membrane
lumenal side of membrane
ER to Golgi transport vesicle membrane
COPII−coated ER to Golgi transport vesicle
MHC class II protein complex
endosome lumen
cytokine receptor binding
cytokine activity
antigen binding
peptide antigen binding
immune receptor activity
MHC protein complex binding
double−stranded RNA binding
MHC class II protein complex binding
NAD+ ADP−ribosyltransferase activity
protein ADP−ribosylase activity
GeneRatio
Count
p.adjust
B)
COiMg - upregulated
immune effector process
regulation of defense response
adaptive immune response
positive regulation of immune response
T cell activation
response to virus
leukocyte mediated immunity
defense response to virus
adaptive immune response based on immunoglobulin recombination
defense response to symbiont
side of membrane
external side of plasma membrane
receptor complex
secretory granule membrane
phagocytic vesicle
MHC protein complex
lumenal side of membrane
integral component of lumenal side of endoplasmic reticulum membrane
lumenal side of endoplasmic reticulum membrane
MHC class II protein complex
immune receptor activity
cytokine receptor binding
cytokine activity
cytokine binding
antigen binding
cytokine receptor activity
peptide antigen binding
MHC protein complex binding
MHC class II protein complex
hydrolase activity, acting binding on carbon−nitrogen (not peptide) bonds, in cyclic amidines
GeneRatio
Count
p.adjust
C)
CO
response to virus
defense response to virus
defense response to symbiont
adaptive immune response
negative regulation of viral process
COiMg
response to virus
defense response to virus
defense response to symbiont
adaptive immune response
immune effector process
-log(q)
D)
CO - downregulated
blood circulation
muscle system process
extracellular matrix organization
extracellular structure organization
external encapsulating structure organization
potassium ion transport
hormone metabolic process
terpenoid metabolic process
membrane repolarization
regulation of cardiac muscle cell membrane repolarization
extracellular matrix
external encapsulating structure
collagen−containing extracellular matrix
collagen trimer
metal ion transmembrane transporter activity
hormone binding
potassium ion transmembrane transporter activity
extracellular matrix structural constituent
peptide hormone binding
voltage−gated sodium channel activity
GeneRatio
Count
p.adjust
E)
COiMg - downregulated
response to xenobiotic stimulus
response to ketone
cell fate commitment
heart morphogenesis
response to corticosteroid
phenol−containing compound metabolic process
catecholamine metabolic process
catechol−containing compound metabolic process
catechol−containing compound biosynthetic process
catecholamine biosynthetic process
extracellular matrix
external encapsulating structure
collagen−containing extracellular matrix
non−motile cilium
platelet alpha granule lumen
G protein−coupled receptor activity
extracellular matrix structural constituent
catecholamine binding
ATPase−coupled inorganic anion transmembrane transporter activity
GeneRatio
Count
p.adjust
F)
CO
blood circulation
extracellular matrix organization
extracellular structure organization
external encapsulating structure organization
membrane repolarization
COiMg
response to xenobiotic stimulus
response to ketone
response to corticosteroid
catechol biosynthetic process
catecholamine biosynthetic process
-log(q)

### Figure S11

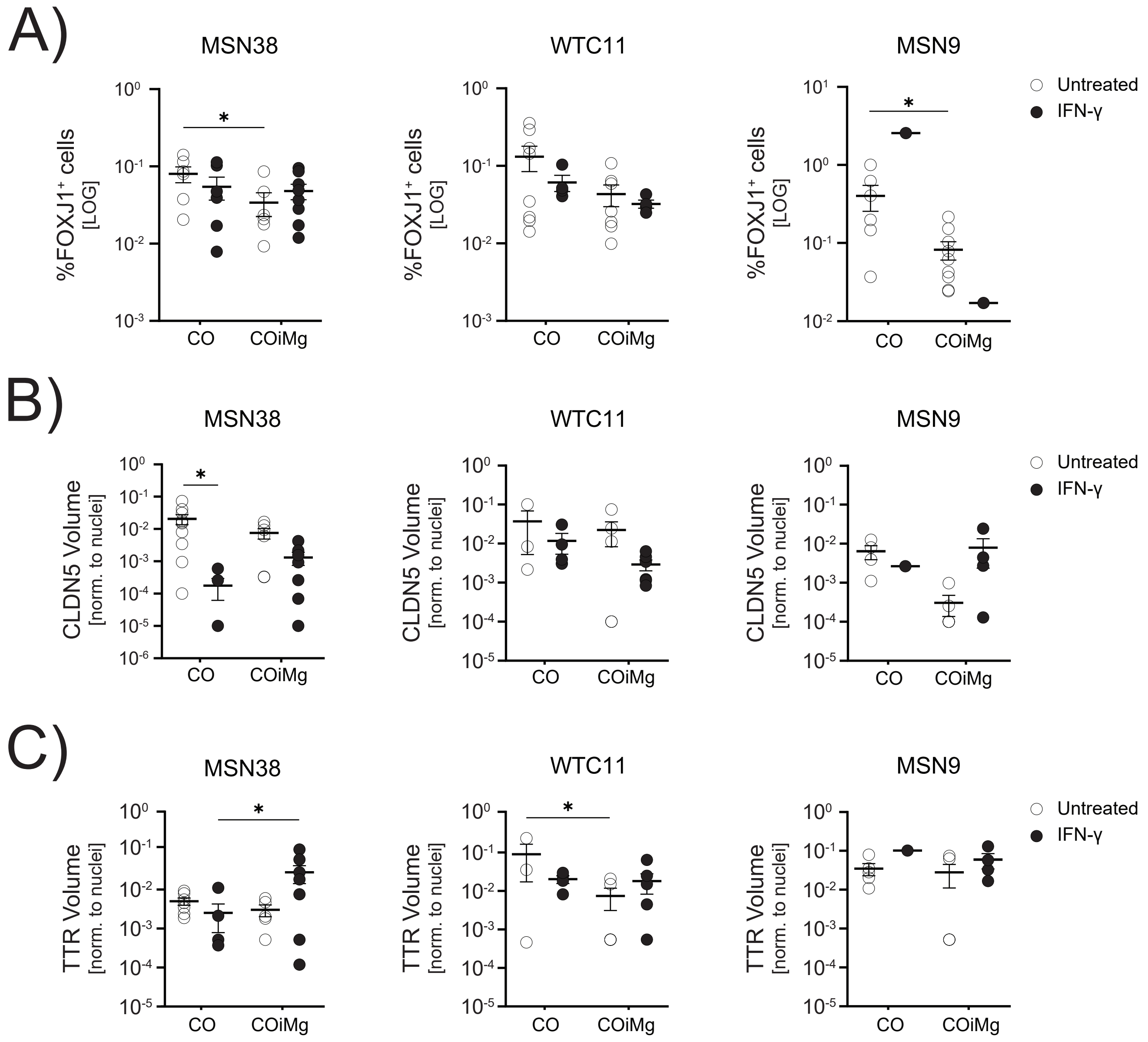

A)
MSN38
WTC11
MSN9
%FOXJ1⁺ cells [LOG]
10^0
10^-1
10^-2
10^-3
10^1
CO
COiMg
*
Untreated
IFN-γ
B)
MSN38
WTC11
MSN9
CLDN5 Volume [norm. to nuclei]
10^0
10^-1
10^-2
10^-3
10^-4
10^-5
10^-6
CO
COiMg
*
Untreated
IFN-γ
C)
MSN38
WTC11
MSN9
TTR Volume [norm. to nuclei]
10^0
10^-1
10^-2
10^-3
10^-4
10^-5
CO
COiMg
*
Untreated
IFN-γ
